# Developmentally regulated *tcf7l2* splice variants mediate transcriptional repressor functions during eye formation

**DOI:** 10.1101/607713

**Authors:** Rodrigo M. Young, Kenneth B. Ewan, Veronica P. Ferrer, Miguel L. Allende, Jasminka Godovac-Zimmermann, Trevor C. Dale, Stephen W. Wilson

## Abstract

Tcf7l2 mediates Wnt/β-Catenin signalling during development and is implicated in cancer and type-2 diabetes. The mechanisms by which Tcf7l2 and Wnt/β-Catenin signalling elicits such a diversity of biological outcomes are poorly understood. Here, we study alternatively spliced *tcf7l2* in zebrafish and show that only splice variants that include exon 5 and an analogous human *tcf7l2* variant can effectively provide compensatory repressor function to restore eye formation in embryos lacking *tcf7l1a/tcf7l1b* function. Knockdown of exon 5 specific *tcf7l2* variants in *tcf7l1a* mutants also compromises eye formation and these variants can effectively repress Wnt pathway activity in reporter assays using Wnt target gene promoters. We show that the repressive activities of exon5-coded variants are likely explained by their interaction with Tle co-repressors. Furthermore, phosphorylated residues in Tcf7l2 coded exon5 facilitate repressor activity. Our studies suggest that developmentally regulated splicing of *tcf7l2* can influence the transcriptional output of the Wnt pathway.

## Introduction

Wnt signalling has a broad array of biological functions, from regional patterning and fate specification during embryonic development to tissue homeostasis and stem cell niche maintenance in adult organs (van Amerongen and Nusse, 2009, Nusse and Clevers, 2017). Because of its relevance to such a diversity of processes, Wnt pathway misregulation is linked to a range of diseases such as cancer and diabetes and neurological/behavioural conditions (Nusse and Clevers, 2017). Wnts can activate several intracellular pathways, and the branch that controls gene expression works specifically through β-catenin and the small family of T-Cell transcription factors (Tcfs; Cadigan and Waterman, 2012).

In absence of Wnt ligand, intracellular β-catenin levels are kept low by a mechanism that involves phosphorylation by GSK-3β and CK1α, which is mediated by the scaffolding of β-catenin by Axin1 and APC in what is termed the destruction complex (MacDonald and He, 2012, Niehrs, 2012). Phosphorylated β-catenin is ubiquitilated and degraded in the proteasome (MacDonald et al., 2009, Niehrs, 2012). In this context, Tcf transcription factors actively repress the transcription of downstream genes by interacting with Gro/TLE like co-repressors (Cadigan and Waterman, 2012, Hoppler and Waterman, 2014). When cells are exposed to Wnt ligand, the destruction complex is disassembled and β-catenin is no longer phosphorylated and committed to degradation (MacDonald et al., 2009, Niehrs, 2012). This promotes the translocation of β-catenin to the nucleus where it displaces co-repressors by its interaction with Tcfs, activating the transcription of Wnt target genes (Cadigan and Waterman, 2012, Hoppler and Waterman, 2014, Schuijers et al., 2014). Hence, Tcf proteins are thought to work as transcriptional switches that can activate transcription in the presence of Wnt ligands or repress transcription in their absence.

During development, ensuring appropriate levels of Wnt/β-catenin signalling is essential for many processes. For instance, during gastrulation, specification of the eyes and telencephalon can only occur when Wnt/β-catenin signalling is low or absent and overactivation of the pathway in the anterior neuroectoderm mispatterns the neural plate leading to embryos with no eyes (Kim et al., 2000, Heisenberg at al., 2001, Kiecker and Niehrs, 2001, Houart et al., 2002, Dorsky et al., 2003). Illustrating this, fish embryos mutant for *axin1*, a member of the β-catenin destruction complex, are eyeless because cells fail to phosphorylate β-catenin, leading to abnormally high levels of the protein, mimicking a Wnt active state (Heisenberg et al, 2001). Similarly, it has been shown that *tcf7l1a/headless* mutants also mimic Wnt/β-catenin overactivation suggesting that it is necessary to actively repress Wnt/β-catenin target genes for regional patterning to occur normally (Kim et al., 2000, Young et al., 2019).

In vertebrates, Lef/Tcf transcription factors constitute a family of four genes: *lef1*, *tcf7 (tcf1)*, *tcf7l1 (tcf3)* and *tcf7l2 (tcf4)*. All contain a highly conserved β-catenin binding domain (β-catenin-BD) at the amino-terminal (N-terminal) end and a high mobility group box (HMG-box) DNA binding domain (DNA-BD) in the middle of the protein (Fig. 1A, Cadigan and Waterman, 2012, Hoppler and Waterman, 2014). All Tcf proteins bind the 5’-CCTTTGATS-3’ (S=G/C) DNA motif, but can also bind to sequences that diverge from this consensus (van de Wetering et al., 1997, van Beest et al., 2000, Hallikas et al., 2006, Atcha et al., 2007). The fact that all Tcfs bind to the same motif has led to the notion that the functional specificity of Lef/Tcf proteins may be imparted by inclusion or exclusion of functional motifs by alternative transcription start sites or alternative splicing (Hoppler and Waterman, 2014).

**Fig. 1.**
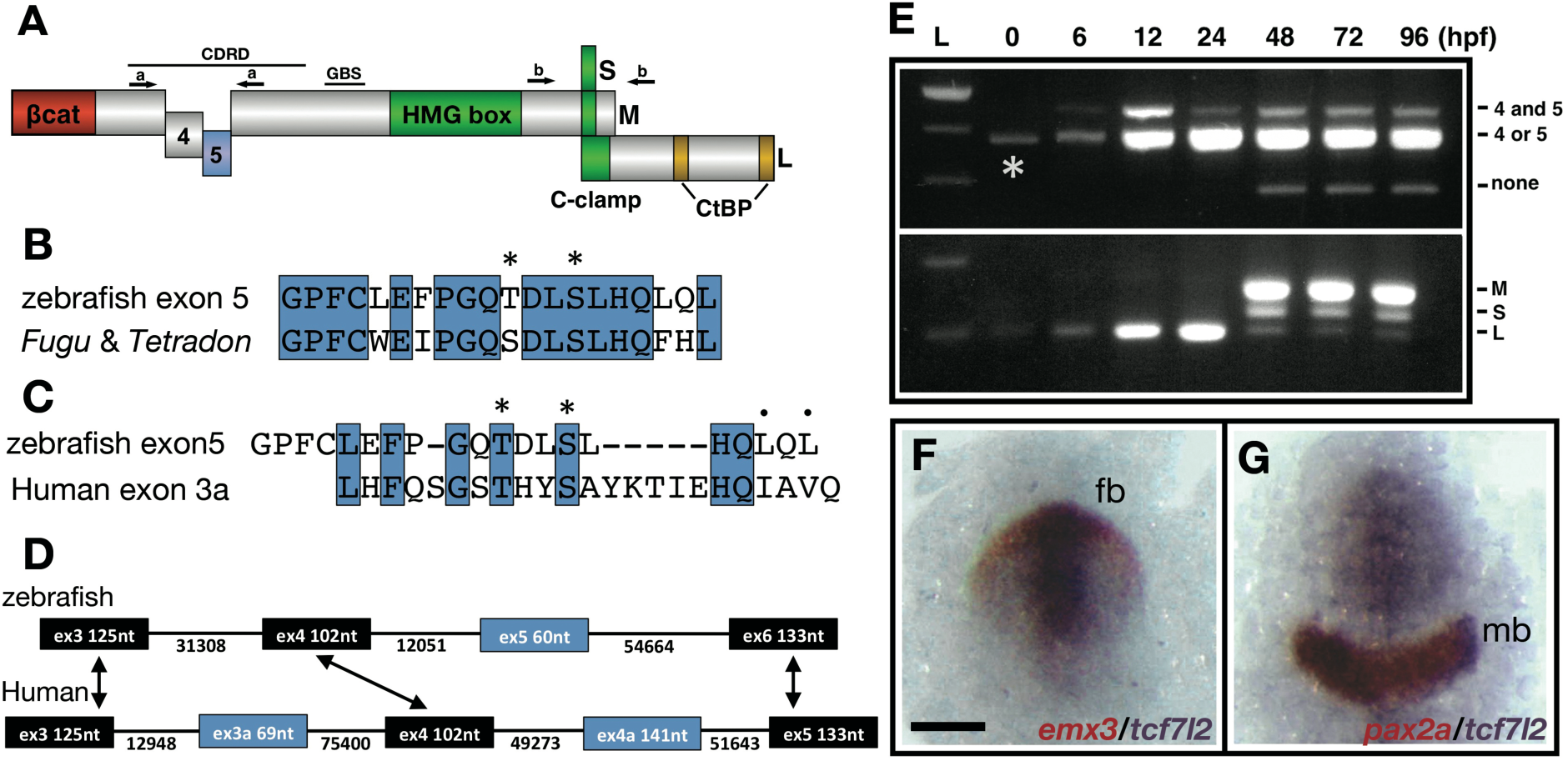
Description and expression of a new alternatively spliced exon in zebrafish *tcf7l2*. **(A)** Schematic representation of variants of Tcf7l2 arising from different splice forms (not to scale). Labels 4 and 5 represent the region of Tcf7l2 coded by alternative exons 4 and 5. Short (S), Medium (M) and Long (L) C-terminal variants coded by alternative splice variants in the 5’ end of exon 15 are indicated. Red box, β-catenin (βcat) binding domain. Green boxes, High-Mobility Group (HMG) Box, which is the primary DNA interacting domain, and C-clamp DNA-helper binding domain. Yellow boxes, CtBP interaction domains. CDRD labelled line over exons 4 and 5 indicates the Context Dependent Regulatory Domain and GBS (Groucho Binding Site) marks the region of interaction with Groucho/Tle transcriptional co-repressors. Arrows indicate the position of primer sets ‘a’ and ‘b’ used for RT-PCR experiments in (E). **(B-C)** Alignment of the amino acid sequences coded by zebrafish, *Takifugu rubripens* and *Tetradon tcf7l2* exon 5 (B) or human exon 3a (C). Identical amino acids marked by blue boxes. Asterisks over sequence mark putative phosphorylated amino acids. Dots over sequence indicate similar amino acids. **(D)** Schematic of the genomic region of zebrafish and human *tcf7l2*. Introns depicted as lines and exons as boxes. Blue exon boxes depict human *tcf7l2* alternative exons 3a and 4a, and zebrafish alternative exon 5. Black exon boxes indicate equivalent exons in both species emphasised by arrows. Numbers under introns and within exons represent their nucleotide size (not to scale). **(E)** RT-PCR experiments performed on cDNA from embryos of ages indicated in hours post fertilisation (hpf). L, 1Kb ladder. Top panel shows results of PCRs using primer set ‘a’ (materials and methods) amplifying the region of alternative exons 4 and 5. Bottom panel shows results of PCRs using primer set ‘b’ (materials and methods) amplifying the region of alternative exon 15. Asterisk shows maternal expression of *tcf7l2*. **(F-G)** Double *in situ* hybridisation of *tcf7l2*, in blue, and *emx3* (F) or *pax2a* (G), in red. 10hpf flat mounted embryos, dorsal view, anterior up, posterior down; fb, prospective forebrain; mb, prospective midbrain. Scale Bar in (F) is 200µm.

The region in between the β-catenin-BD and the DNA-BD of Lef/Tcf proteins, known as the context-dependent regulatory domain (CDRD), and the carboxy-terminal (C-terminal) end of the protein, are coded by alternatively spliced exons (Fig. 1A, Young et al., 2002, Archbold et al., 2012, Cadigan and Waterman, 2012, Hoppler and Waterman, 2014). The C-terminal region of Lef/Tcfs includes the C-Clamp domain and two CtBP interacting motifs (Fig. 1A, Brannon et al., 1999, Valenta et al., 2003, Atcha et al., 2007, Hoverter et al., 2012, Hoverter et al., 2014). In certain contexts, the C-clamp domain helps DNA binding and increases the selectivity for certain gene promoters (Atcha et al., 2007, Wohrle et al., 2007, Chang et al., 2008, Weise et al., 2010). The CDRD includes the domain of interaction with Gro/TLE co-repressors (GBS, Fig. 1A, Cavallo et al., 1998, Roose et al., 1998, Daniels and Weis, 2005, Arce et al., 2009, Chodaparambil et al., 2014) and amino acids that can promote Lef/Tcf dissociation from DNA, modify nuclear localisation or promote activation of transcription when phosphorylated by HIPK2, TNIK or NLK (Shetty et al., 2005, Mahmoudi et al., 2009, Hikasa et al., 2010, Ota et al., 2012). Exons 4 and 5 and the borders of exon7 and exon 9 that are included in the region of the CDRD are alternatively spliced in *tcf7l2* (Duval et al., 2000, Pukrop et al., 2001, Young et al. 2002). The inclusion of the border of exon 9 can transform Tcf7l2 into a strong transcriptional repressor (Liu et al., 2005). Hence, splicing regulation in the CDRD could, similarly, be relevant to transcriptional output (Tsedensodnom et al., 2011, Koga et al., 2012). However, the function of the alternatively spliced exons 4 and 5 is still unknown. All the described variations in Lef/Tcf proteins may contribute to their functional diversity, an idea that is supported by the fact that Tcfs control many different subsets of genes (Cadigan and Waterman, 2012, Hrckulak et al., 2016).

Tcf7l2 has various known roles including a requirement during establishment of left-right asymmetry in the habenulae and for the maintenance of the stem cell compartment in colon and skin epithelia (Korinek et al., 1998, Nguyen et al., 2009, Hüsken et al., 2014). Additionally, polymorphisms in the genomic region that codes for human *tcf7l2* exon 3a (Fig1. C, D), segregate with acquisition of type-2 diabetes (Grant et al., 2006), and conditional knockdowns of *tcf7l2* give rise to mice with phenotypes comparable to diabetic patients (Boj et al., 2012). *tcf7l2* has an alternative translation start site and alternative splicing in the CDRD and in exons that lead to shorter C-terminal ends (Duval et al., 2000, Young et al., 2002, Vacik et al., 2011). Overall, this suggests that many regulatory inputs could influence the transcriptional output of Tcf7l2.

In this study, we address the role of alternative splicing in mediating the functional properties of Tcf7l2 during early nervous system development. Our results show that alternative splicing of *tcf7l2* significantly impacts the transcriptional repressor activity of the encoded protein. *tcf7l2* splice variants have been characterised in humans and, to a lesser extent, in mice and zebrafish (Duval et al., 2000, Young et al., 2002, Prokunina-Olsson et al., 2009, Weise et al., 2010) but little information is available on different roles for the splice variants. In zebrafish, *tcf7l2* is first expressed in the anterior neuroectoderm by the end of gastrulation (Young et al., 2002) and in this study, we show that at this stage, *tcf7l2* is only expressed as long C-terminal variants that can include a newly identified alternative exon 5. We show that only zebrafish Tcf7l2 variants that include the coded exon 5 and comparable human Tcf7l2 variants effectively provide the repressive function required to promote eye specification. Moreover, only these variants are effective in repressing Wnt target gene promoters in luciferase assays, probably due to interaction with Tle co-repressors. We further show that two phosphorylated amino acids coded by exon 5 of Tcf7l2 are required for this interaction, and overall repressive function. Hence, our results suggest that alternative exon 5 in zebrafish *tcf7l2* could play a critical role in mediating transcriptional repression of Wnt pathway target genes. Our data also suggests that through inclusion of the region coded by exon 5, Tcf7l2 could be part of a phosphorylation regulatory module that keeps the Wnt pathway in an ‘off’ state by phosphorylating β-catenin in the cytoplasm and Tcf7l2 in the nucleus.

## Results

### Characterisation of a novel *tcf7l2* alternative splice variant

With the aim of addressing the functional roles of different *tcf7l2* splice variants, we first cloned the zebrafish splice forms in the CDRD (Fig. 1A). This region of Tcf proteins is close to the fragment that interacts with Gro/TLE like co-repressors (GBS, Fig. 1A) and consequently alternative splice forms may affect transcriptional function by modulating interactions with Gro/TLE proteins (Duval et al., 2000, Young et al., 2002, Roose et al., 1998, Daniels and Weis, 2005). Using primers flanking the region containing putative alternative exons in the CDRD encoding region (Primer set-a, Fig. 1A), we performed RT-PCR and cloned the resulting DNA fragments.

The amplified DNA contained a new exon not previously described in zebrafish or in any other species (Fig. 1B, Fig. S1, accession no xxxxx). This exon (*tcf7l2* exon 5 hereafter) codes for a 20 amino acid stretch and is flanked by consensus splice acceptor and donor intron sequences. This region of human *tcf7l2* also includes the alternatively spliced exons 3a and 4a (Fig. 1D, Prokunina-Olsson, 2009). Zebrafish *tcf7l2* exon 5 is similar in size to human *tcf7l2* exon 3a but instead lies in a genomic location that in the human gene would be positioned between exons 4 and 5, where alternative human exon 4a is located (Fig. 1D). Although the protein sequence homology with other fish species is high (Fig. 1B, Fig. S2A), the amino acid identity encoded by human exon 3a and zebrafish exon 5 is only 33% (Fig. 1C). However, in both species, all neighbouring exons are the same size, show a high degree nucleotide and protein sequence homology, and are surrounded by long introns (Fig. 1D, Fig. S2B). Moreover, both fish and human exons 5/3a encode residues that are putative CK1/PKA kinase phosphorylation sites (Fig. 1B, C, asterisks).

RT-PCR experiments showed that splice variants that include alternative exon 4 are expressed maternally and zygotically (Fig 1E, upper panel, middle band labelled ‘4 or 5’, asterisk). Exon 5 is expressed zygotically and is included from 6hpf (hours post fertilisation, Fig. 1E, upper panel, top band labelled ‘4 and 5’). *tcf7l2* splice variants that include exon 5 and lack exon 4 and that lack both exons 4 and 5 (Fig. 1E, upper panel, lower band labelled ‘none’) are only expressed from 48hpf onwards.

The inclusion of two alternative forms of exon 15 adds a premature stop codon that leads to medium and short Tcf7l2 Ct variants (Young et al., 2002). We characterised this region by analysing the alternatively spliced 5’ end of *tcf7l2* exon 15 for the presence of long, medium or short splice variant C-terminal (Ct) coding ends (Fig. 1A,E, lower panel). Maternally, and until 24hpf, *tcf7l2* is only expressed as transcripts that lead to long (L) Tcf7l2 variants (Fig. 1E, bottom panel, lower band). From 48hpf onwards *tcf7l2* is predominantly expressed as splice forms that code for medium (M) and short (S) Ct Tc7f7l2 variants, with L variants barely detectable (Fig. 1E, bottom panel, top two bands).

We further assessed how the expression of exons 4 and 5 relate to alternative splice forms in exon 15 affecting the Ct domain of Tcf7l2. Before 48hpf, *tcf7l2* is only expressed as splice forms that lead to long Ct variants, but from 48hpf, variants including exon 4 are expressed predominantly as medium variants, but also as short and long Ct-forms (Fig. S3A). On the other hand, at 48hpf and onwards, transcripts including exon 5 are expressed only as medium or short Tcf7l2 Ct-variants and by 96hpf only as M variants (Fig. S3B). The range of Tcf7l2 variants expressed as development proceeds is summarised in Table S1A and B.

Given the functional relevance of Tcf7l2 in adult tissue homeostasis (Nusse and Clevers 2017), we studied splicing events involving exons 4/5 and exon 15 in adult zebrafish eye, brain, gut, liver, pancreas, ovaries and testis (Fig. S3C-F). A summary of the RT-PCR results and Tcf7l2 variants expressed in adult organs based on this RT-PCR data is presented in Table S2A and B.

From here, we focus on *tcf7l2* splice variants expressed as the anterior neuroectoderm is patterned during gastrula and early somite stages.

### *tcf7l2* is broadly expressed in the anterior neural plate

The analyses above show that by early somite stage (12hpf), *tcf7l2* is predominantly expressed as two long Ct isoforms, both which include exon 4. One that lacks exon 5 (*4L-tcf7l2* variant from here onwards) is expressed maternally and zygotically, and the other which includes exon 5 is expressed zygotically (*45L-tcf7l2* variants from here onwards).

From late gastrula stage, *tcf7l2* is expressed in the anterior neural plate (Young et al., 2002), overlapping with *tcf7l1a* and *tcf7l1b* (Kim et al., 2000, Dorsky et al., 2003). At 10hpf, the expression of *tcf7l2* overlaps rostrally with that of *emx3* (Fig. 1F, Morita et al., 1995) delimiting the prospective telencephalon, and caudally with the rostral limit of midbrain marker *pax2.1* (Fig. 1G).

Consequently, *tcf7l2* is expressed throughout most of the prospective forebrain including the eye field during the stages when the neural plate becomes regionalised into discrete domains.

### Zebrafish *tcf7l2* exon 5 and human *tcf7l2* exon 3a containing variants are able to restore eye formation upon loss of *tcf7l1a/b* function

Zygotic *tcf7l1a/headless^m881/m881^* mutants (Z*tcf7l1a^-/-^* from here onwards) show reduced eye size at 30hpf but later, through compensatory growth recover eye size (Fig. 2G, Table S3, Young et al., 2019). However, these mutants are sensitised to further loss of Tcf repressor activity and when *tcf7l1b*, the paralogue of *tcf7l1a,* is knocked down by injecting 0.12pmol of a validated ATG morpholino (mo*^tcf7l1b^* Dorsky et al., 2003) in *Ztcf7l1a* mutants, eyes fail to be specified (Fig. 2D, Table S4, 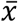=99%, n=270, 3 experiments, Dorsky et al., 2003). To address whether there are any differences in the functional properties of *tcf7l2* splice variants expressed at 12hpf (Fig. 1E, Table S1B), we assessed their ability to restore eye formation when Tcf7l1a/Tcf7l1b function was abrogated (Fig. 2D). Z*tcf7l1a^-/-^* embryos were co-injected with 0.12pmol of mo*^tcf7l1b^* and 20pg of either *4L-tcf7l2* or *45L-tcf7l2* mRNAs. Control overexpression of these *tcf7l2* variants in wildtype or *Ztcf7l1a^+/-^* embryos did not induce any overt phenotype (not shown).

**Fig. 2.**
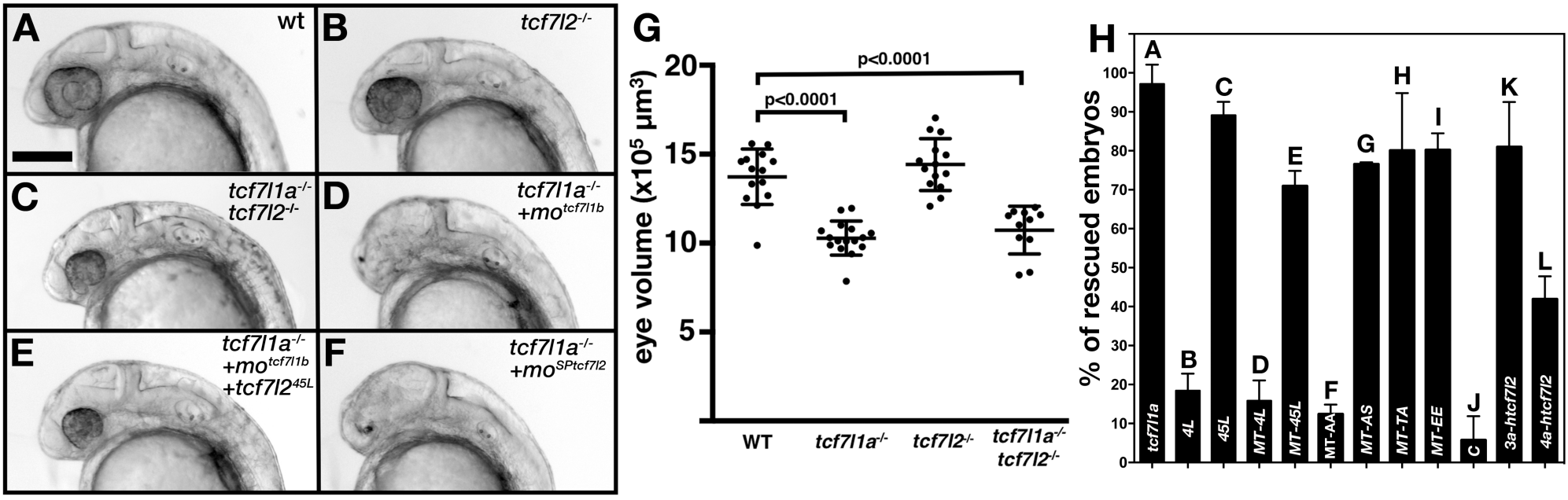
Alternative exon 5 of *tcf7l2* impacts eye formation. **(A-F)** Lateral views (anterior to left, dorsal up) of 28hpf live wildtype (A), *tcf7l2 ^zf55/ zf55^* (B), double *tcf7l1a^-/-^/tcf7l2 ^zf55/ zf55^* (C) and *tcf7l1a*^-*/-*^ (D-F) zebrafish embryos with injected reagents indicated top right showing representative phenotypes. (D) 0.12pmol mo*^tcf7l1b^* (E), 0.12pmol mo*^tcf7l1b^* and 20pg of *45L-tcf72* splice variant mRNA, (F) 1.25pmol mo*^SPtcf7l2^*. Scale bar in (A) is 200µm. **(G)** Plot showing the area of the profile of eyes of 30hpf fixed embryos coming from a double heterozygous *tcf7l1a/tcf7l2* mutant incross. Error bars are mean ±SD, only P values greater than 0.1 from unpaired t test with Welch’s correction are indicated. Data in Table 3. **(H)** Bars represent the percentage of *tcf7l1a^-/-^* embryos that develop eyes (with distinguishable lens and pigmented retina) coming from multiple *tcf7l1a^+/-^* female to *tcf7l1a^-/-^* males crosses, injected with 0.12pmol of mo*^tcf7l1b^* (all bars) and co-injected with constructs stated on X axis: 10pg of *tcf7l1a* mRNA (A), 20pg of *tcf7l2* mRNA splice variants *4L-tcf7l2* (B), *45L-tcf7l2*, (C), *MT-4L-tcf7l2* (D), *MT-45L-tcf7l2* (E), MT-*tcf7l2*-AA (F), MT-*tcf7l2*-AS (G), MT-*tcf7l2*-TA (H), MT-*tcf7l2*-AA (I), *htcf7l2-C* (J), *htcf7l2-3a* (K), and *htcf7l2-4a* (L). Data for all these plots is included in TableS5. Error bars are mean ±SD.

Exogenous *45L-tcf7l2* mRNA effectively rescued the eyeless phenotype in *Ztcf7l1a^-/-^*/*tcf7l1b* morphant embryos (Fig. 2E, H, TableS5, 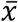=89.1±3.4%, n=163 embryos, 3 experiments), whereas *4L-tcf7l2* mRNA did not (Fig. 2H, TableS5, 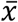=18.4±4.4%, n=146 embryos, 3 experiments). This suggests that only *tcf7l2* splice variants that include exon 5 have the ability to repress the aberrant pathway activation upon loss of Tcf7l1a/Tcf7l1b function.

An alternative explanation for the poor restoration of eye formation by the 4L*-*Tcf7l2 variant could be that it is either not localised to the nucleus or has lower protein stability. To address this, we transfected HEK293 cells with constructs encoding N-terminal Myc tagged (MT) constructs of Tcf7l2. Both MT-Tcf7l2 variants were expressed and localised to the nucleus (Fig. S4), suggesting that the lack of rescue with *4L-tcf7l2* mRNA is due to other differences in protein function. The MT-45L-Tcf7l2 variant was also able to restore eye formation in *Ztcf7l1a^-/-^* mutants injected with mo*^tcf7l1b^* to a similar extent as the untagged variant (Fig. 2H; TableS5, 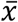 =70.9±3.9%, n=199 embryos, 3 experiments). These results suggest that 45-Tcf7l2 and not 4L-Tcf7l2 variants can compensate for loss of Tcf7l1a/b in eye specification.

To assess the functional activity of human *tcf7l2* alternative exons in the region of zebrafish *tcf7l2* exon5, we cloned long Ct human *tcf7l2* splice variants exclusively including alternative exons 3a (*3a-htcf7l2*) or 4a (*4a-htcf7l2*), or excluding both exons (*C-htcf7l2*). *3a-htcf7l2* variant mRNA was able to restore eye formation in most *Ztcf7l1a^-/-^*/*tcf7l1b* morphant embryos (Fig. 2H, TableS5, 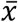=80.9%±11.6, n=? embryos, 3 experiments). However, *4a-htcf7l2* variant mRNA restored eye formation less effectively (Fig. 2H, TableS5, 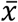=41.9±5.8%, n=? embryos, 3 experiments), and the variant lacking both alternative exons failed to restore eye formation (Fig. 2H, TableS5, 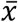=3.6±4.4%, n=? embryos, 3 experiments).

### Eye formation is compromised in *tcf7l1a* mutants when Tcf7l2 lacks the region encoded by exon5

In the *zf55* (*exl*) mutant allele of *tcf7l2,* the first intron of the gene is retained in the mRNA which knocks down expression of the protein generated by the *tcf7l2* transcripts with reading frames starting in exon 1 (Muncan et al., 2007). We found that homozygous *tcf7l2^zf55^* mutants do not show reduced eye size at 30hpf (Fig. 2B, G, Table S3, n=14, data from one of three experiments yielding similar results). To assess if *tcf7l2* repressor activity can functionally compensate for loss of *tcf7l1a,* we incrossed double heterozygous *tcf7l1a^+/m881^/tcf7l2^+/zf55^* fish. However, we did not observe an obvious eye-size phenotype in *Ztcf7l1a/tcf7l2* double homozygous mutants with eye size in these mutants similar to *Ztcf7l1a^-/-^* eyes (Fig. 2C, G. Table S3, three experiments). Neither did injection of *tcf7l2* ATG morpholino in *Ztcf7l1a^-/-^* embryos lead to any enhancement of the eye phenotype (not shown). These results suggest that the lack of severe eye phenotypes in *tcf7l2* and *Ztcf7l1a/tcf7l2* double homozygous mutants is not due to genetic compensation, as has been observed for some other genes (El-Brolosy et al, 2019, Ma et al., 2019).

To further explore if the *45L-tcf7l2* splice variant could play a role in eye formation, we used 1.25pmol/embryo of a splicing morpholino (mo*^SPtcf7l2^*) that targets the intron/exon splice boundary 5’ to exon 5 (Fig. S5A). This Mo is predicted to force the splicing machinery to skip exon5 and splice exon4 to exon6, such that only 4L-Tcf7l2 variants would be translated. The efficacy of the mo*^SPtcf7l2^* was confirmed by RT-PCR, which shows that the band corresponding to *45L-tcf7l2* mRNA is absent in mo*^SPtcf7l2^* injected morphants, but the band corresponding to *4L-tcf7l2* mRNA is still present (Fig. S5B). Sequencing of the putative *4L-tcf7l2* band showed that the exon4/6 splicing event in mo*^SPtcf7l2^* injected morphants is in frame and only leads to the expression of the *4L-tcf7l2* variant (not shown). As expected, Tcf7l2 protein was still present when detected by Western blot (FigS5B). Wildtype embryos injected with mo*^SPtcf7l2^* showed smaller eyes at 32hpf compared to control morpholino (mo^C^) injected embryos (Fig.S5, C-E, n=10, TableS6) and more dramatically, injection of mo*^SPtcf7l2^* in *Ztcf7l1a^-/-^* mutants led to a fully penetrant eyeless phenotype (Fig. 2F, H, Table S4, 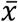=99%, n=128, three experiments). No eyeless phenotype was observed when mo*^SPtcf7l2^* was injected in sibling heterozygous embryos or when a scrambled control morpholino (mo^C^) was injected in *Ztcf7l1a^-/-^* mutants (not shown). These results suggest that specific loss of exon5 coded sequence in Tcf7l2 can compromise its ability to promote eye formation. The strong penetrance of the eyeless phenotype in *Ztcf7l1a^-/-^*/mo*^SPtcf7l2^* is perhaps surprising given that *Ztcf7l1a^-/-^ /tcf7l2^zf55/zf55^* embryos showed no severe eye phenotype, and suggests that an appropriate balance in the levels of *tcf7l2* splice variants may be required for the eye to form normally in *tcf7l1a* mutants.

### Tcf7l2 splice variant with exon 5 shows repressor activity in luciferase reporter assays

Given the diversity of biological outputs driven Wnt/β-catenin signalling and mediated by Tcf transcription factors (Hoppler and Waterman, 2014), the function of Tcf7l2 variants could potentially be to activate/repress the transcription of different subsets of genes. To explore whether 4L and 45L-Tcf7l2 variants show differing promoter transactivation abilities, we performed luciferase assays using the generic TOPflash reporter and known promoters of the Wnt pathway regulated genes, *cdx1* (Hecht and Stemmler, 2003), *engrailed* (McGrew et al., 1999), *cJUN* (Nateri et al., 2005), *lef1* (Hovanes et al., 2001) and *siamois* (Brannon et al., 1999). All the promoters of these genes used in the luciferase assays contain consensus Tcf binding elements.

HEK293 cells were transiently transfected with the luciferase reporter construct and DNA encoding either: 1. the GSK-3β binding domain of mouse Axin2 Flag tag fusion (Flag-Ax2) construct which competes with GSK-3β leading to increased β-catenin levels and Wnt pathway activation (FlagAx-(501-560) in Smalley et al., 1999), or 2. a constitutively-active VP16-TCF7L2 fusion protein, which can induce the expression of Wnt-responsive promoters in absence of nuclear β-catenin (Ewan et al., 2010). As expected, all the tested reporters showed a strong response to Flag-Ax2 or VP16-TCF7L2 transfection (Fig. 3A-F, Table S7 and Table S8, second bar in all plots). We then assessed whether Tcf7l2 splice variants with or without exon 5 could influence the luciferase reporter activation by co-transfecting *Flag-Ax2* (Fig 3A-F, Table S7) or *VP16-TCF7L2* (Fig 3G-L, Table S8) along with either *4L-tcf7l2* or *45L-tcf7l2* splice variants. 45L-Tcf7l2 variant co-transfection led to reduced transactivation by FlagAx2 compared to 4L-Tcf7l2 on all the promotors we tested (Fig 3, Table S7). Moreover, 45L-Tcf7l2 also showed a greater ability to compete with VP16-TCF7L2 when tested with all promotors except for *cjun,* which did not respond to any *tcf7l2* variant co-transfection (Fig. 3G Table S8). Our results suggest that Tcf7l2 variants including exon5 are either less able to activate transcription or are able to repress the transcription at certain promoters.

**Fig. 3.**
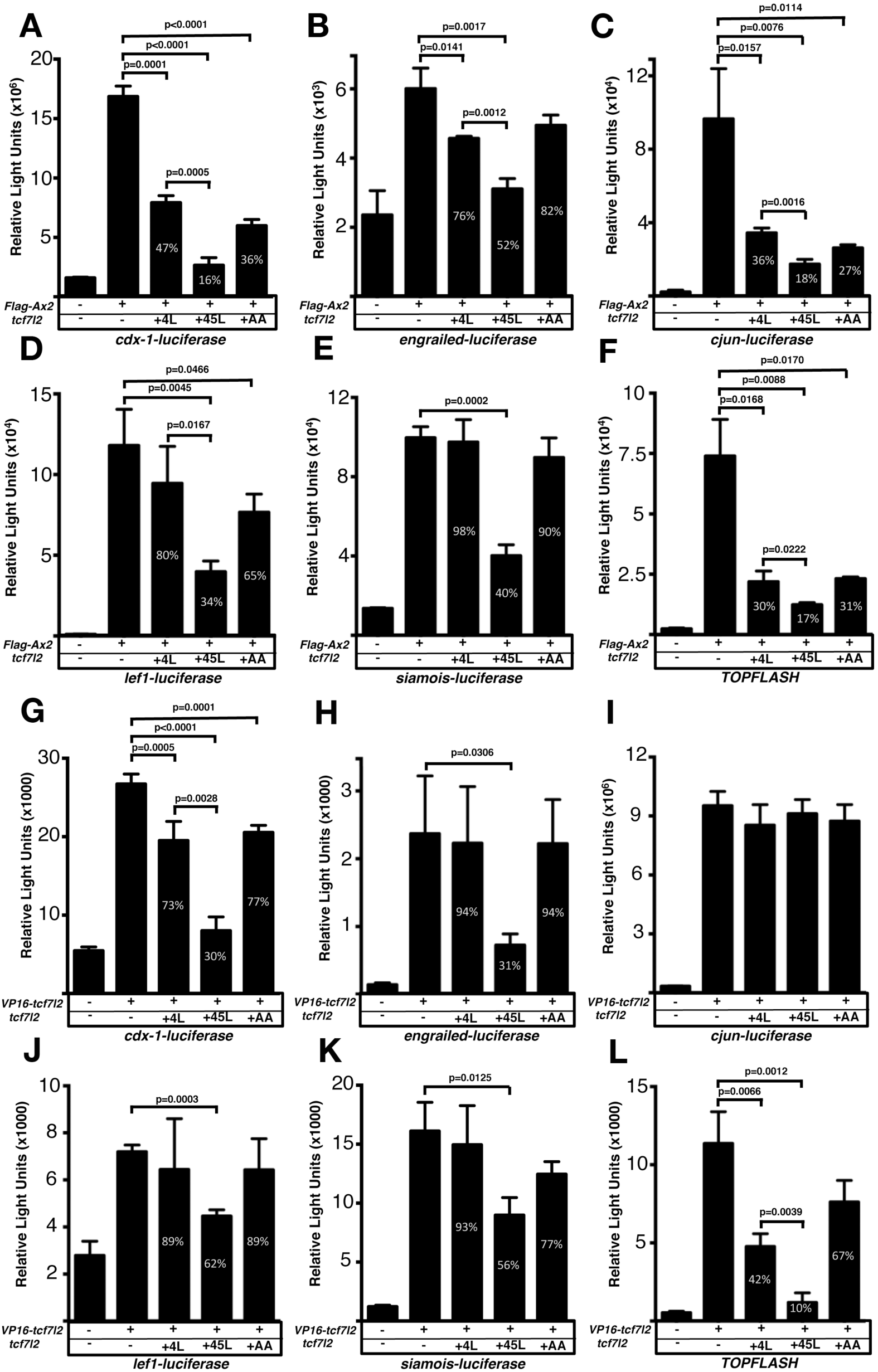
Exon 5 coding *tcf7l2* variant repress Wnt activity induced by *FLAG-Ax2* or *VP16-TCF7L2* in luciferase assays. Bar plots showing luciferase reporter assay results expressed in relative light units. HEK293 cells were transiently co-transfected with luciferase reporter constructs indicated beneath the X-axis, (**A-F**) *FLAG-Ax2* (except for first bars), (**G-L**) *VP16-TCF7L2* DNA (except for first bars) and *4L-tcf7l2* DNA (+4L; 3^rd^ bars), or *45L-tcf7l2* DNA (+45L; 4^th^ bars) or *tcf7l2-AA* DNA (+AA; 5^th^ bars). Control experiments show only background luciferase activity with no transfected plasmids (1^st^ bars). Figures in the bars indicate the percentage size of that bar relative to transfection with either *FLAG-Ax2* or *VP16-TCF7L2* alone (2^nd^ bar). Error bars are mean ±SD n=3 (experiments were performed twice), P values from unpaired t tests comparing *FLAG-Ax2* or *VP16-TCF7L2* control condition with *tcf7l2* variant co-transfections. Comparisons with no statistical significance are not marked.

### Inclusion of exon 5 enhances the interaction between Tcf7l2 and Tle3b

The region of Tcf7l2 encoded by exon 5 is located in the vicinity where Lef/Tcf proteins interact with Gro/TLE-like corepressors (Roose et al., 1998, Brantjes at el., 2001, Daniels and Weis, 2005, Arce et al., 2009). Consequently, inclusion of exon 5 in *tcf7l2* mRNA could potentially assign repressor activity by modifying the interaction of Tcf7l2 with Gro/TLE proteins. Alternatively, inclusion of exon 5 could modify the capacity of Tcf7l2 to interact with transactivating β-catenin.

To address if the inclusion of exon5 can modulate the interaction between Tcf7l2 and Gro/TLE or β-catenin proteins, we performed yeast two-hybrid (Y2H) protein interaction experiments between 4L-Tcf7l2 and 45L-Tcf7l2 variants and Tle3b (the zebrafish orthologue of mammalian Tle3/Gro1), or β-catenin (Fig. S6). Full-length Tle3b seemed to be either toxic or have transfection problems in the yeast strain and so we used a C-terminal deletion of Tle3b (dC-Tle3b), which still included the glutamine-rich domain that interacts with Tcf proteins (Daniels and Weis, 2005). Yeast co-transfected with β*-catenin* or *tle3b* and *4L-tcf7l2* or *45L-tcf7l2* splice variants were able to grow in complete auxotrophic selective media (-L-A-H-W + Aureoblastinin) and also express the X-gal selection reporter (Fig. S6). This suggests that both Tcf7l2 variants are able to interact with β-catenin and dC-Tle3b. However, this Y2H assay cannot reveal differences in the affinity of interactions between proteins.

To address possible differences in protein interactions between Tcf7l2 splice variants and β-catenin or Tle3b, we performed co-immunoprecipitation (co-IP) experiments using protein extracts from co-transfected HEK293 cells. Both Myc tagged Tcf7l2 variants were efficiently immunoprecipitated by anti-Myc beads (Fig. 4, right top panel) and showed a similar capacity to co-IP with β-catenin (Fig. 4, right middle panel, second and third lanes). However, MT-45L-Tcf7l2 showed a significantly higher capacity to co-IP Tle3b compared to 4L-Tcf7l2 (Fig. 4, right bottom panel, second and third lane, asterisk). This suggests that the repressor activity of MT-45L-Tcf7l2 observed in our *in vivo* and luciferase reporter assays could be mediated by an enhanced interaction with Gro/TLE-like co-repressors.

**Fig. 4.**
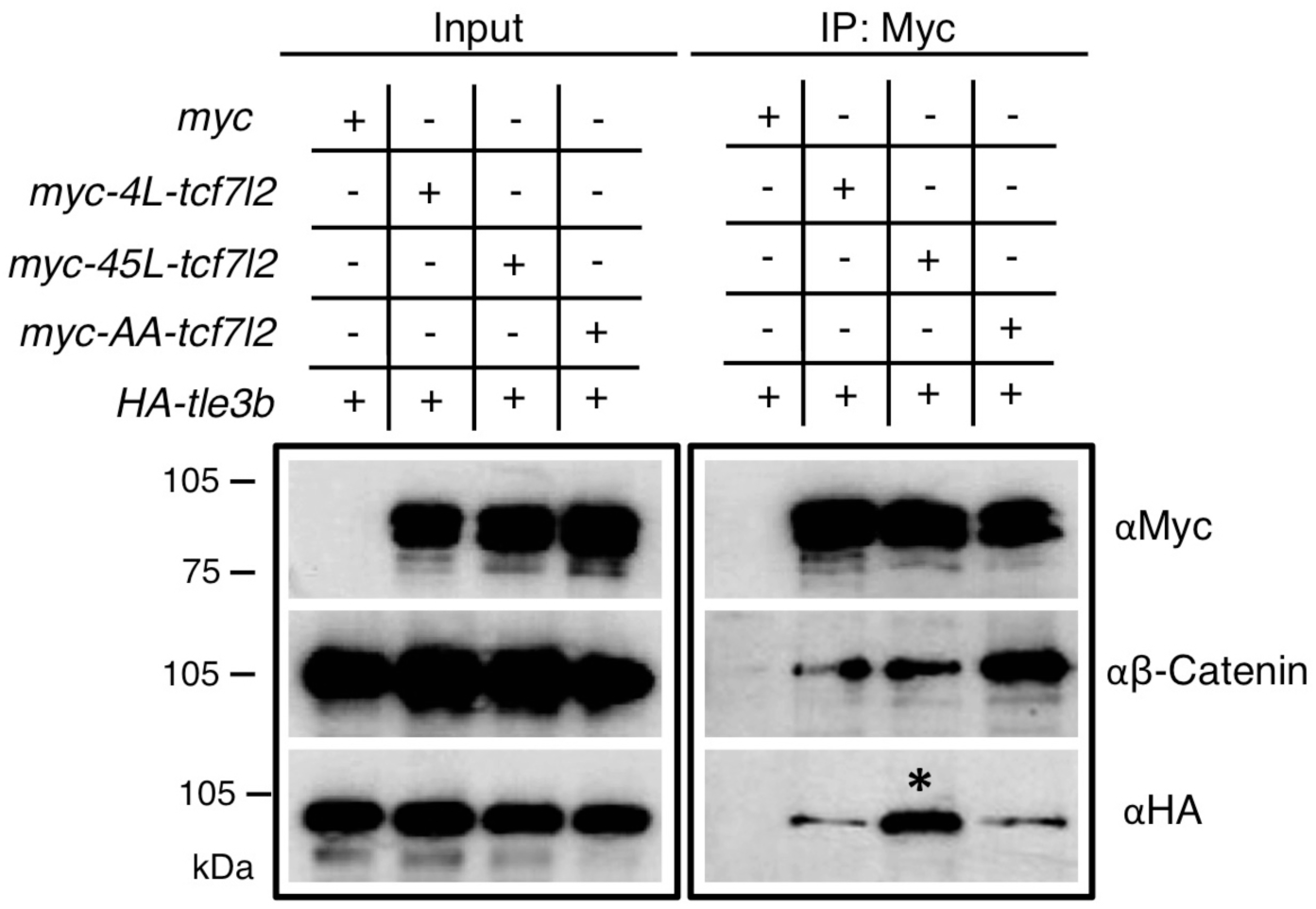
Alternative exon 5 of *tcf7l2* enhances affinity with Tle3b. Protein input (left panel) and anti-Myc immunoprecipitation (IP) eluate western blot (right panel) showing co-inmunoprecipitation of β-Catenin or HA-tagged Tle3b. HEK293 cells were transiently transfected with HA tagged *tle3b* together with empty myc tag vector (1^st^ lane), *MT*-*4L-tcf7l2* (2^nd^ lane), *MT-45-tcf7l2* (3^rd^ lane) and *MT-AA-tcf7l2* (4^th^ lane). Left panels show protein input before anti-Myc IP. Right panels show protein eluate from anti-Myc antibody coupled beads. Westernblots were probed with anti-Myc (tagged Tcf7l2 proteins, top panel), anti-βcatenin (middle panel) and anti-HA (tagged Tle3b protein, bottom panel) antibodies. Asterisk shows that the Tcf7l2 form containing exon 5 shows more intense binding with Tle3b than other Tcf7l2 forms.

### Phosphorylated amino acids in the domain of Tcf7l2 coded by exon 5 mediate transcriptional repressor function

Kinases are known to modulate Wnt signalling activity through phosphorylation of LRP6, β-catenin and Tcfs (Li et al., 2002, Davidson et al., 2005, Zeng et al., 2005, Sokol, 2011) and bioinformatic analysis using Netphos3.1 (cbs.dtu.dk/services/NetPhos, Blom et al., 2004) or GPS2.0 (<u>gps.biocuckoo.org</u>, Xue et al., 2008) predict that Tcf7l2 exon 5 coded Thr^172^ and Ser^175^ are putatively phosphorylated by GSK-3, CK1, PKA, and other kinases. GSK-3 and CK1 are known to regulate Wnt signalling by phosphorylating LRP6 co-receptor and β-catenin (Niehrs, 2012).

To address if amino acids in Tcf7l2 exon are phosphorylated, we performed mass spectrometry (MS) analysis. N-terminal *MT-4L-tcf7l2* and *MT-45L-tcf7l2* variants were cloned in frame to BioID2 (Kim et al., 2016), an optimised biotin ligase protein, and were transiently expressed in HEK cells treated with biotin. Steptavidin pulled down proteins were used for LC-MS/MS analysis recovering 40% of Tcf712 amino acid sequence (Fig.S7). The recovered peptide containing exon 5-coded sequence, Asp^151^-Lys^184^ was phosphorylated (TableS9).

To address the role of the putatively phosphorylated exon 5 coded amino acids in Tcf7l2, we generated an MT-45L-Tcf7l2 mutant version in which both Thr^172^ and Ser^175^ were replaced by alanines (MT-45L-Tcf7l2-AA), which cannot be phosphorylated. Unlike MT-*45L-tcf7l2*, expression of MT-*45L-tcf7l2-AA* (20pg mRNA per embryo) was unable to rescue the eyeless phenotype of *Ztcf7l1a^-/-^/tcf7l1b* morphant embryos (Fig. 2H, Table S5, 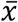=12.4±2.5%, n=117, three experiments). However, *MT-45L-tcf7l2* mutant forms containing Thr172Ala (20pg MT-*45L-tcf7l2-AS* mRNA per embryo, Fig. 2H, Table S5, 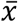=76.6±0.5%, n=34, two experiments) or Ser175Ala (20pg MT-*45L-tcf7l2-TA* mRNA per embryo, Fig. 2H, Table S5, 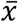=77.9±17.9%, n=44, two experiments) were able to rescue the eyeless phenotype suggesting that repressor activity could be elicited through phosphorylation of either or both of amino acids Thr^172^ and Ser^175^. Furthermore, we generated a Thr172Glu/Ser175Glu phospho-mimicking MT-45L-Tcf7l1a mutant form (MT-45L-Tcf7l2-EE). Expression of *MT-45L-tcf7l2-EE* (20pg mRNA per embryo) restored eye formation in 80.3±4.5% of *Ztcf7l1a^-/-^/tcf7l1b* morphant embryos (Fig. 2H, Table S5, n=48, three experiments).

Moreover, MT-45L-Tcf7l2-AA behaved like MT-4L-Tcf7l2 variants in luciferase reporter assays (Fig. 3, TableS7 and 8, fifth bar in all plots), and also showed a reduced capacity to interact with Tle3b in co-IP experiments (Fig. 4, right bottom panel, fourth lane). Of note, the MT-Tcf7l2-AA mutant showed greater capacity to co-IP with β-catenin compared to MT-4L-Tcf7l2 or MT-45L-Tcf7l2 variants (Fig. 4, right top panel, fourth lane). These results suggest that phospho Thr^172^ and phospho Ser^175^ residues may regulate the ability of 45L-Tcf7l2 to repress target genes and consequently contribute, together with Tcf7l1a/Tcf7l1b function, to zebrafish eye specification.

## Discussion

In this study, we show that developmentally regulated splicing contributes to Wnt signalling regulation during patterning of the anterior neural plate and eyes. We show that there is extensive variety in splicing of zebrafish *tcf7l2* throughout development and across adult tissues and that the exon 5 coded region within Tcf7l2 influences its transcriptional repressor activity. Tcf repressor function is required for maintaining low levels of pathway activity during forebrain and eye specification. We show that long carboxy-terminal Tcf7l2 variants that include both exons 4 and 5 can restore eye formation in fish with compromised *tcf7l1a/b* function and that specific knockdown of the 45L-Tcf7l2 variant in *tcf7l1a* mutants leads to embryos with no eyes. A similar specific human Tcf7l2 variant that includes the region coded by the alternative exon 3a is also able to restore eye formation in *tcf7l1a/b* knockdown embryos, suggesting conservation in the role of splicing in mediating transcriptional repressor activity of Tcf7l2 proteins. Additionally, 45L-Tcf7l2 variants have less capacity to transactivate known Wnt target gene promoters in luciferase reporter assays and can out-compete constitutively transcriptionally active *VP16-hTcf7l2* chimeric proteins. Mass spectrometry results also suggest that the protein region coded by *tcf7l2* exon 5 is phosphorylated and the transcriptional repressor function of the 45L-Tcf7l2 variant requires this phosphorylation potentially due to more efficient interaction with Tle3b corepressor.

### Tcf7l2 transcriptional repression function contributes to forebrain patterning

Like *tcf7l1a* and *tcf7l1b*, *tcf7l2* is expressed in the anterior neural ectoderm, from where the eyes and forebrain develop (Kim et al., 2000, Young et al., 2002, Dorsky et al., 2003). Experimental manipulations or mutations that increase Wnt/β-catenin activity in the anterior neural plate during gastrulation generate embryos with no forebrain and eyes (Wilson and Houart, 2004). For instance, *tcf7l1a* is cell-autonomously required for eye field specification and zebrafish embryos lacking zygotic *tcf7l1a/headless^m881^* function have smaller eyes (Kim et al., 2000, Young et al., 2019). However, this phenotype is exacerbated when *tcf7l1b* is knocked down in *tcf7l1a* mutants leading to embryos with no eyes (Dorsky et al., 2003). The notion that active repression by Tcf7l1a/b transcription factors is required for neuroectodermal patterning is supported by the observation that overexpression of transcriptional dominant active *VP16-tcf7l1a* chimera also leads to eyeless embryos (Kim et al., 2000).

Eyes are smaller in wildtype embryos and absent in in *Ztcf7l1a^-/-^* embryos in which *tcf7l2* variants including exon5 have been knocked down. As only the 45L-Tcf7l2 variant, which includes exon5, can efficiently restore eye formation in *Ztcf7l1a^-/-^/tcf7l1b* morphant embryos, our results suggest that the exon5 coding region can assign transcriptional repressor activity to Tcf7l2 and, as for Tcf7l1 proteins, this transcriptional repressive function contributes to forebrain patterning and eye formation. This conclusion is supported by 45L-Tcf7l2 variants having less transactivation capacity and greater repressor activity compared to 4L-Tcf7l2 and Tcf7l2-AA variants in luciferase reporter experiments; additionally, 45L-Tcf7l2 variants show greater association for Tle3b transcriptional co-repressor in co-IP experiments.

The lack of severe forebrain and eye phenotypes in single *tcf7l1* mutants is at least in part due to the overlapping functional activities of different *tcf* genes and we assume the same is likely for the *tcf7l2* mutant. However, we find no evidence that mutant RNA triggered genetic compensation (el-Brolosy et al. 2019; Ma et al. 2019; El-Brolosy and Stainier 2017; Rossi et al. 2015) is the reason that *tcf7l2* mutants lack severe phenotypes. For instance, morpholino knockdown of *tcf7l2* translation, which circumvents this mechanism, does not result in a severe eye phenotype. Given that, as for other Tcf genes (Cadigan, 2012; Ramakrishnan et al., 2018), *tcf7l2* specific splice variants seem to encode proteins with either repressor or activator roles, mutations that disrupt the balance between these functions may lead to phenotypes differing from complete gene abrogation. Potentially, mis-regulation of repressor function while maintaining the ability of the protein to activate transcription (as suggested by our experiments that specifically knock down *45L-tcf7l2* variants with a splicing morpholino) might lead to forebrain and eye phenotypes that differ from complete loss of protein expression. Hence, the balance between the levels of Tcf transcriptional repressing *versus* activating variants may facilitate correct subdivision of the telencephalic, eye and diencephalic territories of the neural plate.

### Phosphorylation of Tcf7l2 mediates its function as a transcriptional repressor

The context dependent regulatory domain of Tcfs, which includes the region coded by *tcf7l2* exon 5, is acetylated, phosphorylated and sumoylated (Yamamoto et al., 2003, Shetty et al., 2005, Mahmoudi et al., 2009, Hikasa et al., 2010, Ota et al., 2012, Elfert et al., 2013). The importance of phosphorylation to the repressor activity of 45L-Tcf7l2 is suggested by the finding that the phosphorylation resistant Tcf7l2-AA mutant form in which both Thr^172^ and Ser^175^ amino acids are replaced by alanine, behaves as if lacking the region coded by exon 5 in all the functional assays we tested. Conversely, the phospho-mimicking EE-Tcf7l2 variant was able to restore eye development in *Ztcf7l1a^-/-^/tcf7l1b* morphant embryos supporting the idea that phosphorylation of either Thr^172^ or Ser^175^ is required for the repressive function of 45L-Tcf7l2 variants. Although our mass spectrometry analysis confirmed that the peptide encoded by zebrafish *tcf7l2* exon 5 is phosphorylated, the resolution was not sufficient to discriminate whether the phosphorylation was on Thr^172^, Ser^175^ or both. However, mRNA overexpression of both AS-Tcf7l2 and TA-Tcf7l2 mutant variants restores eye formation in *Ztcf7l1a^-/-^/tcf7l1b* morphant embryos, suggesting that the phosphorylation of either amino acid is sufficient to enable the repressor activity of 45L-Tcf7l2.

Gro/TLE transcription co-repressors are displaced by β-catenin to allow Tcf-mediated transcriptional activation (Daniels and Weis 2005, Arce et al., 2009, Chodaparambil et al., 2014) and although we find 45L-Tcf7l2 still interacts with β-catenin, the Tcf7l2-AA mutant variants show greater binding capacity. This raises the possibility that 45L-Tcf7l2 variants may co-exist as phosphorylated and un-phosphorylated pools with different repressor/activator activity.

It is widely accepted that Tcfs work as transcriptional switches, repressing transcription of downstream genes in absence of Wnt ligand, and activating gene transcription when Wnt signalling is active (Cadigan, 2012; Ramakrishnan et al., 2018). In this context, the repressive function of Tcf7l2 may be part of an integrated pathway response when the Wnt pathway is not active. The ‘off’ state of Wnt signalling involves active phosphorylation of β-catenin by CK1α and GSK-3β kinases (MacDonald and He, 2012, Niehrs, 2012). Our findings support a model in which Tcf transcription factors could also be part of a kinase regulatory module that maintains the pathway in an ‘off’ state, not only in the cytoplasm by phosphorylating β-catenin, but also by promoting transcriptional repression through phosphorylation of Tcf7l2. Direct assessment of an *in vivo* functional role for phosphorylation of Tcf7l2, and potentially other Tcfs, will be required to address such a model.

The salience of resolving a role for phosphorylation in the regulation of transcriptional activation/repression is heightened given the relevance of Tcf7l2 in mediating colorectal cancer outcome due to imbalanced Wnt signalling. Indeed, one future avenue for investigation will be to identify proteins that interact with the phosphorylated and un-phosphorylated forms of Tcf7l2.

### Functional modulation of Tcf7l2 through spatial and temporal regulation of splicing of alternative exons

The occurrence of widespread tissue specific and developmentally regulated *tcf7l2* splicing suggests that certain variants of Tcf7l2 are required for proper cell and tissue type specification during development and for its various roles in organ physiology during adult life (Nusse and Clevers, 2017). For instance, we have previously shown that Tcf7l2 is required for the development of left-right asymmetry of habenular neurons (Hüsken et al., 2014) and at the stages during which habenular neurons are becoming lateralised, *tcf7l2* splice variants transition from expressing long to only medium and short carboxy-terminal end variants. This is potentially significant as only long variants include a complete C-clamp DNA binding-helper domain (Young et al., 2002) and this domain can direct Tcf7l2 to specific promoters (Hoverter et al., 2014). Consequently, absence of a whole C-clamp may bias the promoter occupancy of Tcf7l2 and shift the expression profile of Wnt target genes (Atcha et al., 2003, 2007, Hecht and Stemmler, 2003, Hoveter et al., 2014).

Tcf7l2 is linked to type-2 diabetes outcome (Grant et al., 2006, Lyssenko et al., 2007, Prokunina-Olsson et al., 2009, Savic et al., 2011) and the strongest risk factor SNPs are located in introns near human *tcf7l2* exon 3a (Grant et al., 2006). Perhaps surprisingly, only liver tissue-specific *tcf7l2* knockouts in mice and no other organs (including pancreas), lead to metabolic outcomes mimicking type-2 diabetes (Boj et al., 2012 and see Bailey et al., 2015). The adult zebrafish liver only expresses 4L and 45L Tcf7l2 variants. Although homologies between fish and human exons in this region of Tcf7l2 are uncertain and despite lack of overall sequence conservation, human *tcf7l2* exon 3a and fish exon 5 have a similar size and share amino acids that are likely phosphorylated in zebrafish Tcf7l2 coded exon 5. Moreover, human *tcf7l2* variants including alternative exon 3a, and to a lesser extent exon 4, can restore eye formation in *Ztcf7l1a^-/-^/tcf7l1b* knockdown embryos. This suggests that the region coded by human Tcf7l2 exon 3a, and possibly exon 4a, may have similar functions as zebrafish exon 5 and that alternative exons in this region may play an evolutionarily conserved role in the transcriptionally repressive capacity of Tcf7l2. Consequently, it will be interesting to assess if the expression levels of these human *TCF7L2* exons is altered in type-2 diabetes patients carrying risk factor SNPs.

Given the importance of maintaining balanced Wnt/β-catenin pathway activity throughout development and tissue homeostasis, elucidating all mechanisms that impact Wnt signalling modulation is critical if we are to understand, and develop ways to manipulate, pathway activity when mis-regulated in pathological conditions. Our work adds weight to the idea that regulation of alternative splicing and controlling the balance between repressor and activator functions of Tcf proteins play an important role in Wnt/β-catenin pathway regulation.

## Materials and Methods

### Animal use, mutant and transgene alleles, genotyping and quantification of eye size

Adult zebrafish were kept under standard husbandry conditions and embryos obtained by natural spawning. Wildtype and mutant embryos were raised at 28°C and staged according to Kimmel et al. (1995). Fish lines used were *tcf7l1a^m881^* (Kim et al., 2000), *tcf7l1b^zf157tg^* (Gribble et al., 2009) and *tcf7l2^zf55^* (Muncan et al., 2007). These three lines are likely to abrogate expression of proteins coded by the reading frame starting in exon 1. There is an alternative downstream transcription start site in mouse *tcf7l2* (Vacik et al., 2011) and likely in other *tcf* genes too (unpublished observations). It is not known if transcripts from these alternative start sites have any functional roles in embryos carrying the mutations above.

Genomic DNA was isolated by HotSHOT method (Suppl. Materials and methods) and *tcf7l1a^m881^* and *tcf7l2^zf55^* mutations were genotyped by KASP assays (K Biosciences, assay barcode 1145062619) using 1µl of genomic DNA for 8µl of reaction volume PCR as described by K Biosciences. Adult zebrafish organs were dissected as described in Supp. Materials and methods. The sizes of eye profiles were quantified from lateral view images of PFA fixed embryos by delineating the eye using Adobe Photoshop CS5 magic wand tool and measuring the area of pixels included in the delineated region. The surface area was then transformed from px^2^ to µm^2^ and then to predicted eye volume as in Young et al., 2019. Embryos were scored as eyeless when no retinal tissue was observed. Even though in the rescue experiments there was variability in the size of the restored eyes, for the sake of simplicity, all embryos with distinguishable eyes were scored as having eyes. This binary categorisation made the rescued versus not-rescued eye phenotype more straightforward to score and less affected by subjective interpretation.

### RNA extraction, reverse transcription and PCR

Total RNA was extracted from live embryos and adult zebrafish using Trizol (Invitrogen) and homogenised by pestle crushing and vortexing. SuperscriptII (Invitrogen) was used for reverse transcription under manufacturers’ instructions using oligo dT and 1µg of RNA for 20µg reaction volume. The following primers were used to amplify fragments of *tcf7l2* cDNA: region exon4/5 (Set a-F TCAAAACAGCTCTTCGGATTCCGAG, Set a-R CTGTAGGTGATCAGAGGTGTGAG), region exon15 (Set b-F GATCTGAGCGCCCCAAAGA AGTG Set b-R CGGGGAGGGAGAAATCATGGA GG).

### mRNA synthesis, embryo microinjection and morpholinos

*tcf7l2* splice variant PCR fragments were cloned in pCS2+ or pCS2+MT expression vectors for mRNA synthesis. mRNA for overexpression was synthesised using SP6 RNA mMessage mMachine transcription kit (Ambion). One to two cell stage embryos were co-injected with 10nl of 5pg of GFP mRNA and morpholinos or *in vitro* synthesised mRNA at the indicated concentrations. Only embryos with an even distribution of GFP fluorescence were used for experiments. Morpholino sequences:

mo*^tcf7l2ATG^* (5’-CATTTTTCCCGAGGAGCGCTAATTT-3’). Embryos injected with this morpholino fail to produce Tcf7l2 protein (Fig. S5).
mo*^SPtcf7l2^* (5’-GCCCCTGCAAGGCAAAGACGGACGT-3’). This splice-blocking morpholino leads to exon skipping to generate a Tcf7l2 protein lacking exon 5 derived amino acids (Fig. S5). *tcf7l1a^-/-^* embryos injected with mo*^SPtcf7l2^* lack eyes, a phenotype not seen when the morpholino is injected into *tcf7l1a^+/-^* siblings. No equivalent genetic mutation that leads to loss of exon 5 of *tcf7l2* exists but the loss of eye phenotype is consistent with other conditions in which the overall level of TCF-mediated repression is reduced (Dorsky et al., 2003).
moC (5’-CTGAACAGGCGGCAGGCGATCCACA-3’). This morpholino is a sequence scrambled version of mo*^SPtcf7l2^*used as an injection control.
mo*^tcf7l1b^* (5’-CATGTTTAACGTTACGGGCTTGTCT-3’; Dorsky et al., 2002)*. tcf7l1a^m881/m881^* embryos injected with mo*^tcf7l1b^* phenocopy the loss of eye phenotype seen in *tcf7l1a^m881/m881^/tcf7l1b^zf157tg/zf157tg^* double mutants (Young and Wilson, unpublished).

### In situ hybridisation and probe synthesis

Digoxigenin (DIG) and fluorescein (FLU)-labelled RNA probes were synthesized using T7 or T3 RNA polymerases (Promega) according to manufacturers’ instructions and supplied with DIG or FLU labelled UTP (Roche). Probes were detected with anti-DIG-AP (1:5000, Roche) or anti-FLU-AP (1:10000, Roche) antibodies and NBT/BCIP (Roche) or INT/BCIP (Roche) substrates according to standard protocols (Thisse and Thisse, 2008).

### Luciferase reporter experiments

The following reporters were used: *cdx1*:luc (Hecht and Stemmler, 2003), *engrailed*:luc (McGrew et al., 1999), *cJun*:luc (Nateri et al., 2005), *lef1*:luc (Hovanes et al., 2001), *siamois*:luc (Brannon et al., 1997), and TOPflash (Molenaar et al., 1996). HEK cells were transfected according to standard methods and using the conditions described in Supp. Materials and Methods.

### Zebrafish protein extraction

Embryos were washed once with chilled Ringers solution, de-yolked by passing through a narrow Pasteur pipette, washed three times in chilled Ringers solution supplemented with PMF (300mM) and EDTA (0.1mM). Samples were briefly spun down, media removed, Laemlli buffer 1X was added at 10µl per embryo and incubated for 10min at 100°C with occasional vigorous vortexing before chilling on ice. Samples were loaded in polyacrylamide gels or stored at −20°C.

### HEK cell transfection, immunohistochemistry and co-immunoprecipitation

HEK cells were grown in 6 well plates and transfected with 4µg of each DNA with lipofectamine 2000 (Invitrogen) for 6hrs according to manufacturers instructions.

For immunohistochemistry, cells were fixed 48hrs after transfection in 4% paraformaldehyde in PBS for 20 min at room temperature, washed with PBS, and permeabilized in 0.2% Triton X-100 in PBS for 5 min at room temperature. The protocol was followed as in Supp. Materials and Methods. For immunoprecipitation, cells were grown for 24hrs after transfection and then proteins were extracted following standard methods (Supp. Materials and Methods). The eluate from antibody beads (30µl) was loaded in 10% polyacrylamide gels and proteins were detected by Western blots (standard conditions) using anti-myc (1/20,000, SC-40, SCBT), anti-HA (1/10,000, 3F10, Roche) and anti-β-catenin (1/8000, Sigma, C7207), to detect the co-immunoprecipitated proteins. Antibodies used on Western blots in Fig. S4 are anti human Tcf7l2 (N-20, SCBT) and anti-gamma tubulin (T9026, Sigma) HRP coupled secondary antibodies (1/2,000, sigma) were used and blots were developed using an ECL kit (Promega).

### Mass Spectrometry experiments

Cell extract proteins were pulled down with streptavidin coated magnetic beads. The protein eluate was run on an SDS-PAGE gel and stained with Coomassie blue. Stained gels were cut, de-stained and Trypsin Gold, Mass Spectrometry Grade (Promega, Madison, USA) in 50 mM ammonium bicarbonate was added in each well containing dried gel pieces and incubated overnight at 37°C. Next day, 0.1% formic acid was added to stop the trypsinolysis and the eluted tryptic peptides were collected in MS glass vials, vacuum dried and dissolved in 0.1% formic acid for LC-MS/MS.

LC-MS/MS analysis was performed with an LTQ-Velos mass spectrometer (Thermo Fisher Scientific, U.K.). Peptide samples were loaded using a Nanoacquity UPLC (Waters, U.K.) with Symmetry C18 180umX20mm (Waters part number 186006527) trapping column for desalting and then introduced into the MS via a fused silica capillary column (100 μm i.d.; 360 μm o.d.; 15 cm length; 5 μm C18 particles, Nikkyo Technos CO, Tokyo, Japan) and a nanoelectrospray ion source at a flow rate at 0.42 μl/min. The mobile phase comprised H_2_O with 0.1% formic acid (Buffer A) and 100% acetonitrile with 0.1% formic acid (Buffer B). The gradient ranged from 1% to 30% buffer B in 95 min followed by 30% to 60% B in 15 min and a step gradient to 80% B for 5 min with a flow of 0.42μl/min. The full scan precursor MS spectra (400-1600 m/z) were acquired in the Velos-Orbitrap analyzer with a resolution of r = 60,000. This was followed by data dependent MS/MS fragmentation in centroid mode of the most intense ion from the survey scan using collision induced dissociation (CID) in the linear ion trap: normalized collision energy 35%, activation Q 0.25; electrospray voltage 1.4 kV; capillary temperature 200 °C: isolation width 2.00. The targeted ions were dynamically excluded for 30s and this MS/MS scan event was repeated for the top 20 peaks in the MS survey scan. Singly charged ions were excluded from the MS/MS analysis and XCalibur software version 2.0.7 (Thermo Fisher Scientific, U.K.) was used for data acquisition. Raw data were analysed using Proteome Discoverer (PD v1.3) with Mascot search engine and Swiss-Prot human and Zebrafish proteome database. Up to 2 trypsin missed cleavages were allowed, carbamidomethylation was set as a fixed modification, while methionine oxidation, phosphorylation of serine, threonine and tyrosine were set as variable modifications. Mass tolerance was set to 8 ppm for the precursors and to 0.6 Da for the fragments.

#### Yeast two-hybrid assays

N-terminal deletions of the first 53 amino acids of *tcf7l2* splice variants were cloned in *pGBK*. Full-length β-catenin and a C-terminal deletion of *tle3b* (NM_131780, complete reading frame after amino acid 210) were cloned in *pGAD* (Clontech). Combinations of plasmids to test two-hybrid interactions were co-transformed in Y2Gold yeast strain (Suppl. Materials and Methods). Transformed yeast were plated on -Leu-Trp dropout selective media agar plates supplemented with X-gal. Positive blue colonies were streaked to an -Ade-His-Leu-Trp dropout selective media agar plates supplemented with Aureoblastidin A and X-gal (Clontech yeast two-hybrid manual).

## Acknowledgments

We thank members of our lab, Florencia Cavodeassi and Mate Varga for stimulating discussions, the UCL fish facility team for fish care and Elke Ober, Richard Dorsky, Marian Waterman, Randy Moon and others for reagents; Masa Kai for advice on Western blot methods and Abdol Nateri for advice on HEK cell protein extraction. We also thank Lucia di Vagno, Graham W Taylor and Mark Crawford for mass spectrometry analysis. This study was generously supported by the MRC (MR/L003775/1 to SW and Gaia Gestri) and Wellcome Trust (088175, 104682/Z/14/Z S.W. and R.Y.), a Marie Curie Incoming International Fellowship (R.Y.), a Royal Society International Joint Project (S.W. and M.A.) and FONDAP (15090007) to (M.A).

## Author contributions

RMY and SWW conceived the study; RMY, KBE and VPF performed experiments; JG-Z performed the mass spectrometry analysis; RMY and SWW wrote the paper with input from all authors.

---

**Fig. S1.**
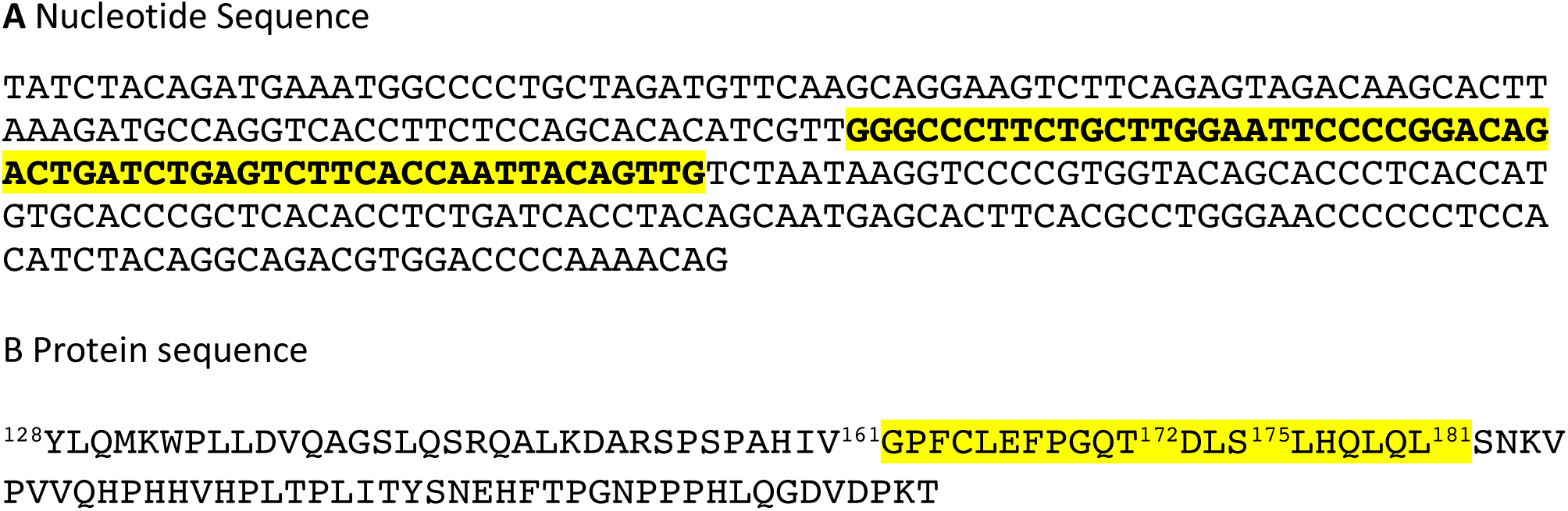
Zebrafish exon 5 nucleotide and coded amino acid sequences. **(A)** Nucleotide sequence of zebrafish *tcf7l2* exon 5 (bold and highlighted) and neighbouring exons. **(B)** Amino acid sequence of the translated sequence of exons in (A).

**Fig. S2.**
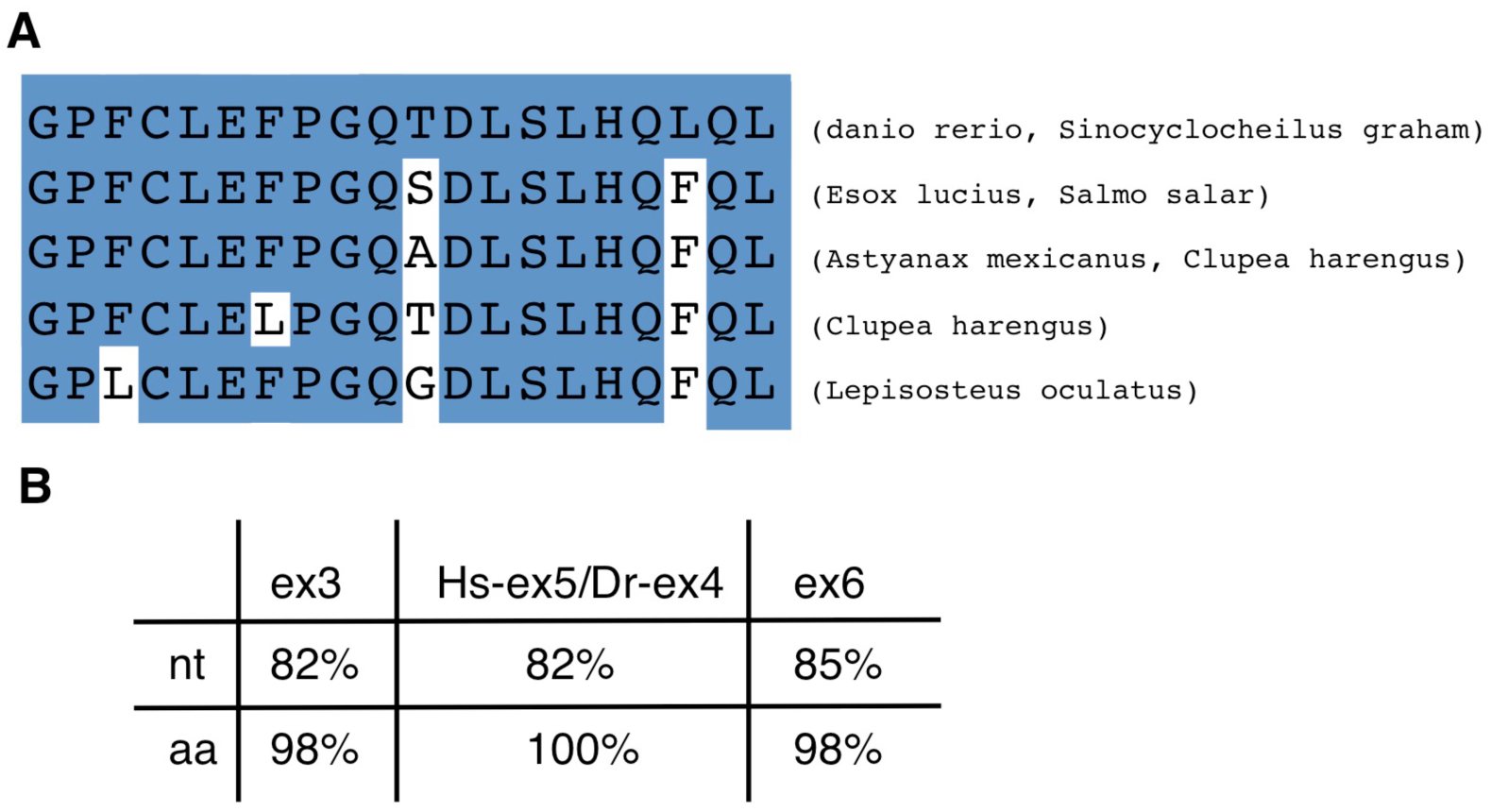
Alignment of the amino acid sequence coded by *tcf7l2* exon 5 in zebrafish and other fish species. **(A)** Alignment of the zebrafish and other fish species amino acid sequence coded by *tcf7l2* exon 5. **(B)** Table showing the nucleotide (nt) homology between human and zebrafish exons surrounding zebrafish new exon 5 and the amino acid (aa) identity of the protein regions they code. Hs, *Homo sapiens*; Dr, *Danio rerio*.

**Fig. S3.**
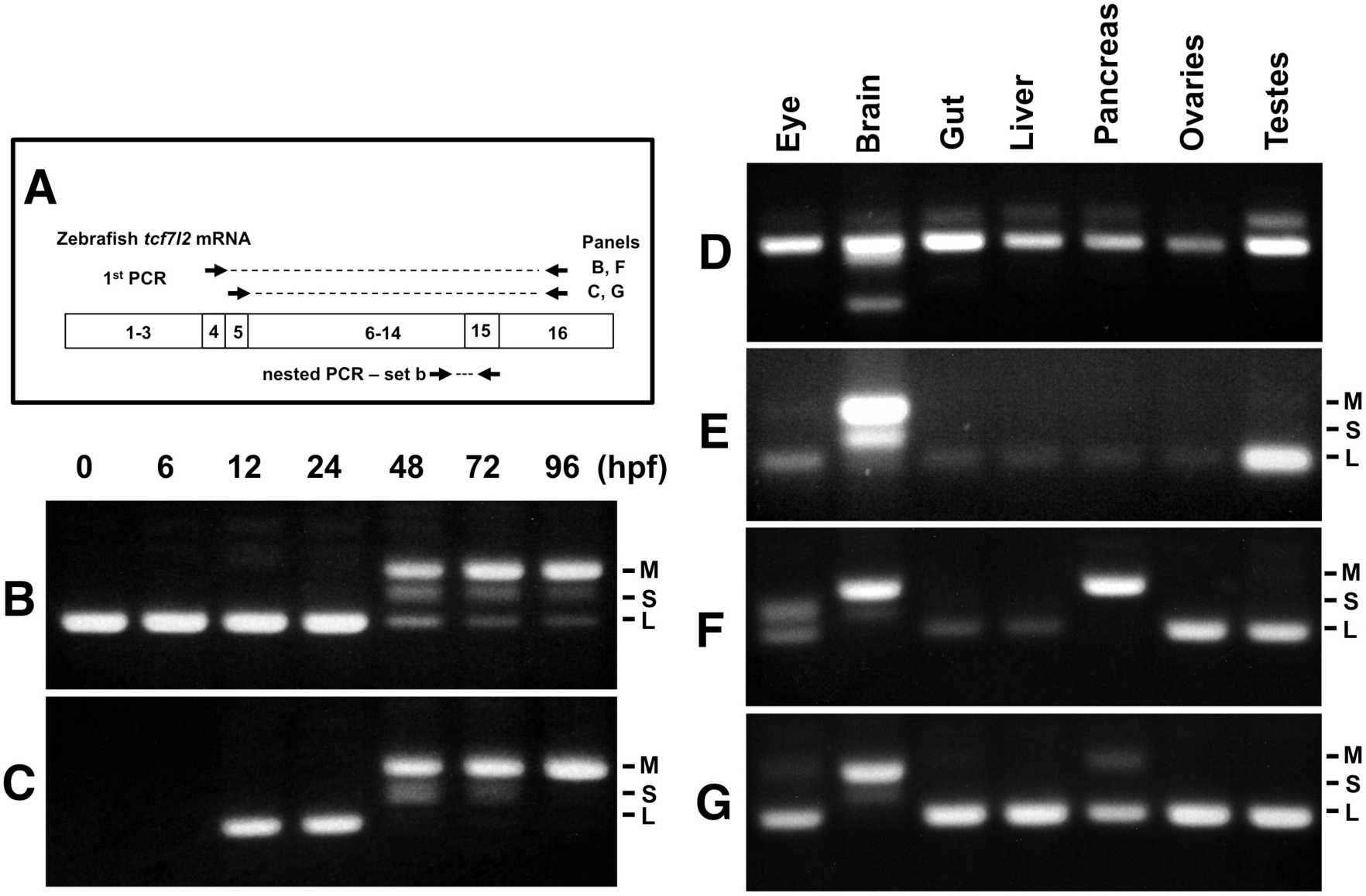
Expression of *tcf7l2* alternative exons 4, 5 and 15 varies across development and in adult organs. RT-PCR analyses of exons 4, 5 and 15 alternative borders of zebrafish *tcf7l2* across development and in various adult organs. **(A)** Schematic representation of nested RT-PCR strategy used for panels (B, C, F and G). **(B-C)** cDNA from embryos of ages indicated (hpf) was PCR amplified using a forward primer that anneals over exon 4 (**B)** or exon 5 (**C)** and a reverse primer that anneals to exon 16 common to all *tcf7l2* mRNA variants. The product from this first PCR was then used as a template for a nested PCR using primer set ‘b’ (as in Fig. 1.A) that amplifies exon 15 and reveals the different Ct ends of Tcf7l2. This last PCR product is shown in these panels. M (Medium), S (short) and L (Long) Ct Tcf7l2 variants. **(D-E)** RT-PCR experiments performed on cDNA of the indicated adult organs using primer set ‘a’ (materials and methods) amplifying the region of alternative exons 4 and 5 (**D**) or using primer set ‘b’ (materials and methods) amplifying the region of alternative exon 15 (**E**). **(F-G)** Same PCR amplification strategy used in panels (**B-C**) to detect the Ct *tcf7l2* variants associated with exons 4 or 5, but using the indicated adult organ cDNA as template in the 1^st^ PCR reaction.

**Fig. S4.**
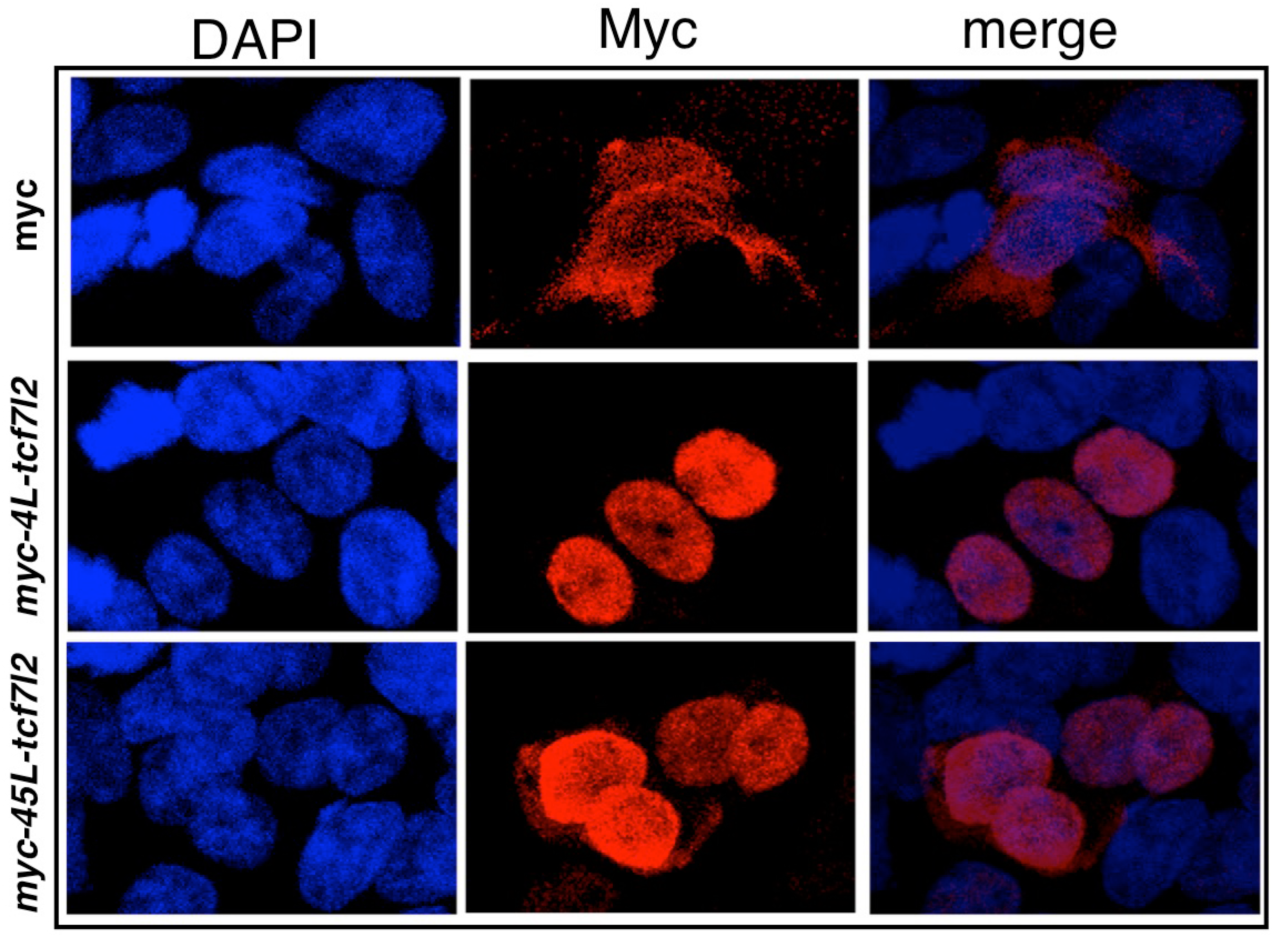
Tcf7l2 variants localise to the nucleus. Sub-cellular localisation of 4L-Tcf7l2 and 45L-Tcf7l2 myc tagged splice variants. HEK293 cells were transfected with empty myc tag vector (top row), *MT-4L-tcf7l2* (middle row) and *MT-45L-tcf7l2* (right row) splice variants and immunostained with anti-myc antibody (2nd column) and stained with DAPI (1st column). Merged images in 3rd column.

**Fig. S5.**
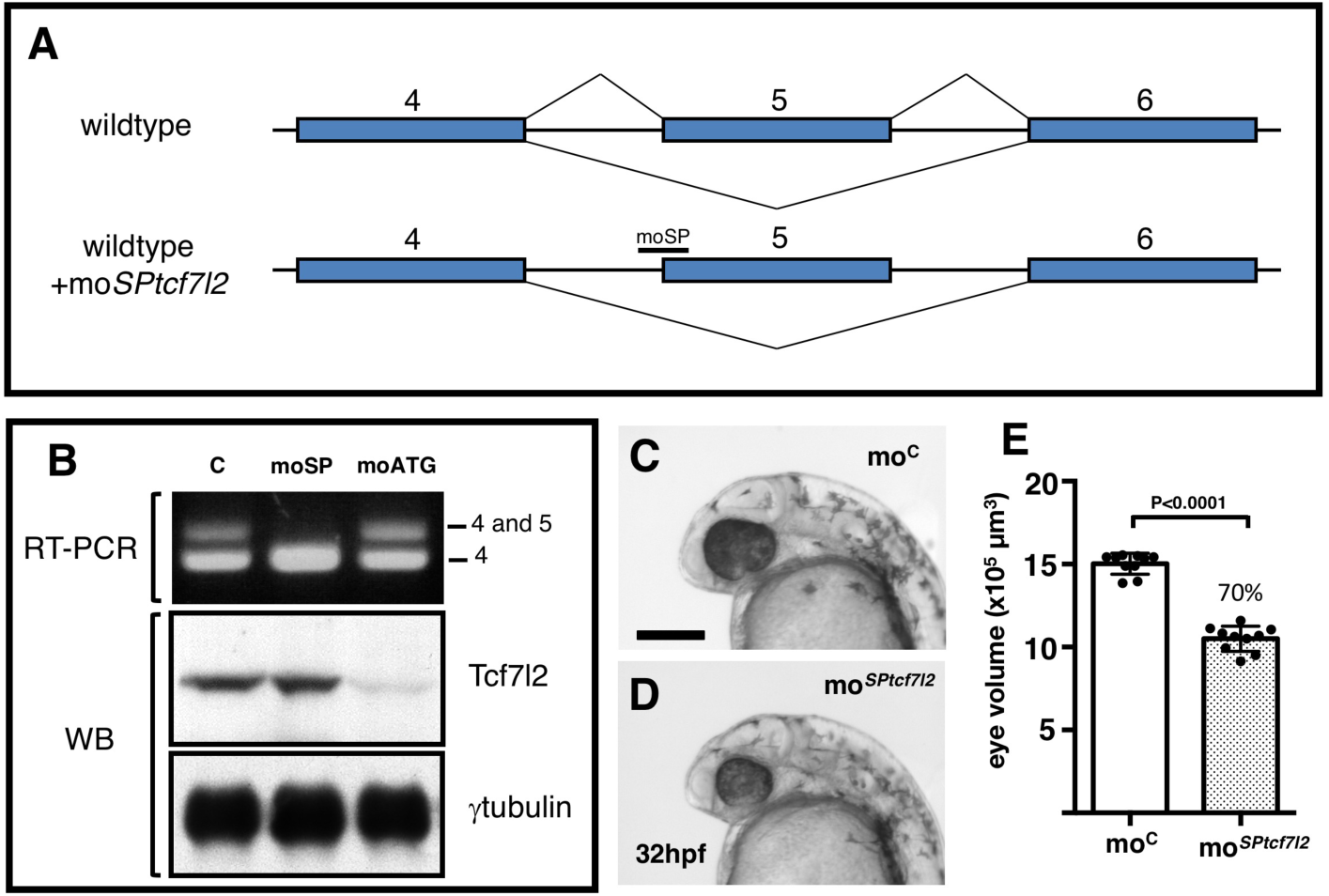
Splicing specific morpholino knockdown of *tcf7l2* splice variants that include exon5. **(A)** Cartoon showing the rationale of using mo*^SPtcf7l2^* that targets intron 4/exon 5 splice site boundary blocking the splicing machinery and making it skip to the next splicing acceptor site in exon 6. **(B)** RT-PCR (**top panel**) and Western blot (**middle and bottom panels**) performed using RNA and proteins extracted from 24hpf zebrafish embryos injected with: control morpholino (first lane), mo*^SPtcf7l2^* (moSP, second lane), and mo*^ATGtcf7l2^* (moATG, third lane). cDNA was amplified using set ‘a’ primers (Fig.1A, 5’F1 vs Splice R1, materials and methods). Anti human Tcf7l2 and anti-gamma tubulin antibodies were used in Western blot experiments (middle and bottom panel). **(C, D**) 32hpf embryos injected with 1.25pmol of (**C**) control morpholino (mo^C^) or (**D**) *tcf7l2* exon 5 splicing morpholino (mo*^SPtcf7l2^*). Scalebar in (**C**) is 200µm. **(E)** Plot showing estimated eye volume of 32hpf live embryos injected with injected with mo^C^ or mo*^SPtcf7l2^* as in (**C, D**). Error bars are mean ±SD, P values from unpaired t test. Percentage indicates average eye profile size relative to control.

**Fig. S6.**
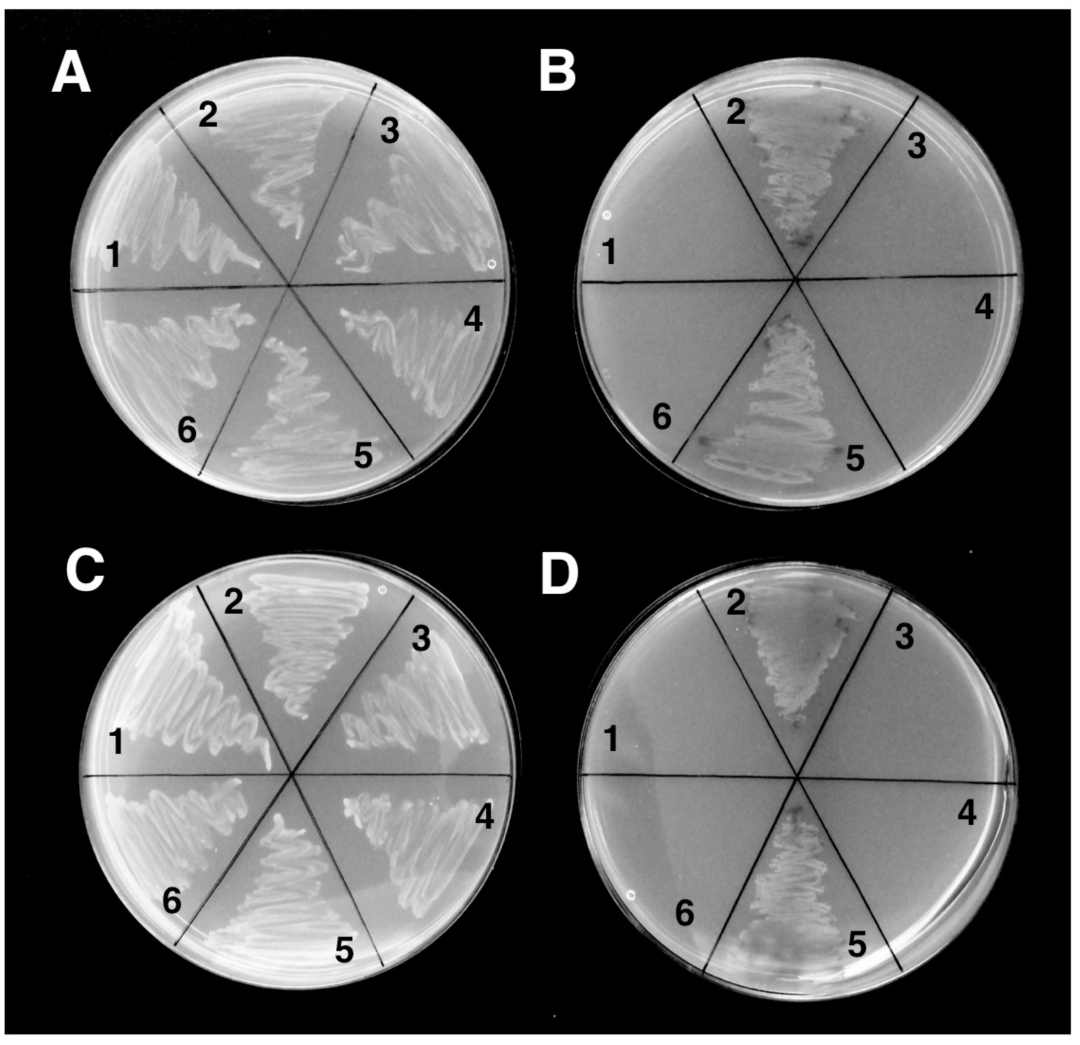
4L-Tcf7l2 and 45L-Tcf7l2 variants interact with β-Catenin and Tle3b in yeast two-hybrid protein interaction assays. Y2Gold yeast strain was co-transformed with: (**1**) β-catenin/*pGAD* and empty *pGBK* vector, (**2**) β-catenin/*pGAD* and *4L-tcf7l2/pGBK* (**A, B**) or *45L-tcf7l2/pGBK* (**C, D**), (**3**) empty *pGAD* vector and *4L-tcf7l2/pGBK* (**A, B**) or *45L-tcf7l2/pGBK* (**C, D**) with, (**4**) *dCtle3b/pGAD* and Empty *pGBK* vector, (**5**) *dCtle3b*/*pGAD* and *4L-tcf7l2/pGBK* (**A, B**) or *45L-tcf7l2/pGBK* (**C, D**), (**6**) empty *pGBK* and *pGAD* vectors. and plated in -Leu-Trp dropout selective media agar plates supplemented with Xgal (**A, C**) or -Ade-His-Leu-Trp dropout selective media agar plates supplemented with Aureoblastidin A and X-gal (**B, D**).

**Fig S7.**
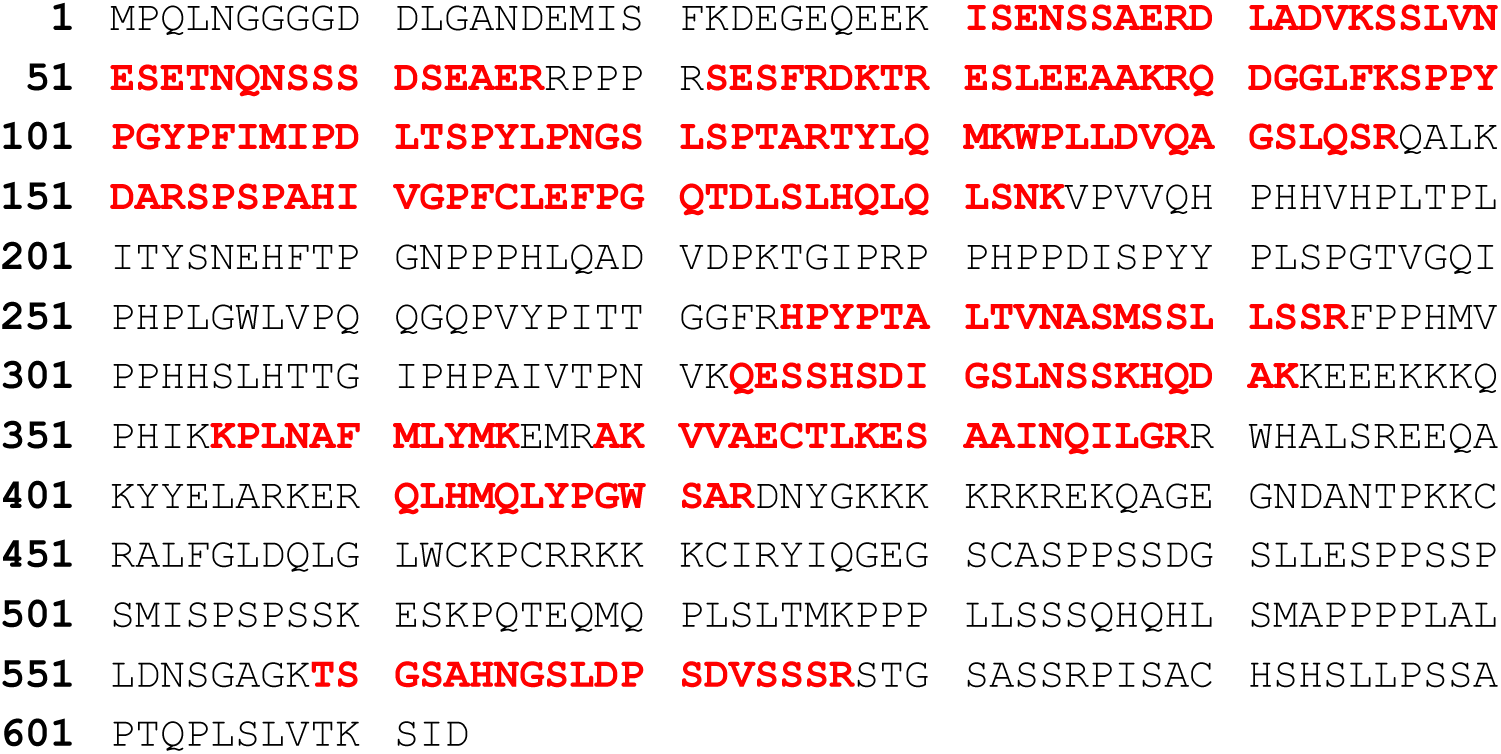
Zebrafish MT-45L-Tcf7l2-BioID2 peptides recovered by LC-MS/MS analyses are shown in bold.

**Table S1 Summary table of RT-PCR analyses showing developmental expression of alternative *tcf7l2* exons 4, 5 and 15 (A) and Tcf7l2 variants (B) through development.**

**(A)** Results in rows 1 to 4 (grey) taken from Fig 1E, rows 5 to 7 (pink) from FigS3A and rows 8 to 10 (blue) from FigS3B. (+) and (++) depict relative band intensity in the gels; (-) indicates no signal

**(B)** Tcf7l2 variants expressed during development based on the information in (A). (+) and (-) indicate presence or absence of the variant respectively. (?) indicates that it is not possible to derive a conclusion based on the data available. Analysis of variants that lack both exons 4 and 5 is not included.

**Table S2 Summary table of RT-PCR analysis of alternative *tcf7l2* exons 4, 5 and 15 (A) and Tcf7l2 variants (B) expressed in adult organs.**

**(A)** Results in rows 1 to 4 (grey) taken from FigS3C, rows 5 to 7 (pink) from FigS3E and rows 8 to 10 (blue) from FigS3F. (+) and (++) depict relative band intensity in the gels; (-) indicates no signal

**(B)** Tcf7l2 variants expressed in adult organs based on the information in (A). (+) and (-) indicate presence or absence of the variant respectively. (?) indicates that it is not possible to derive a conclusion based on the data. Analysis of variants that lack both exons 4 and 5 is not included.

**Table S3 Size of the *tcf7l2^-/-^*eye is similar to wildtype embryos at 30hpf.**

Estimated volume in µm^3^ of the eye of 30hpf fixed embryos coming from a double heterozygous *tcf7l1a/tcf7l2* mutant incross. Avg, average; SD, Standard Deviation.

**Table S4 Knockdown of tcf7l1b and excision of tcf7l2 exon5 in tcf7l1a^-/-^ mutant embryo compromises eye formation**

Embryos from female *tcf7l1a^+/-^* to male *tcf7l1a^-/-^* spawnings were injected with the morpholinos stated in the left column. This pairing scheme leads to 50% of homozygous mutant embryos. Each row represents an individual experiment. Embryos were scored as eyeless when little or no pigmented retinal tissue could be distinguished. Total represents the number of embryos scored in each experiment.

**Table S5. Restoration of eye formation by expression of exogenous Tcf7l2 variants in tcf7l1a^-/-^ /tcf7l1b morphant embryos.**

*Tcf7l1a^-/-^* embryos injected with *tcf7l1b* morpholino and *tcf7l1a* or the *tcf7l2* mRNA variant stated in the first column. Each row represents an individual experiment. Total represents the number of *tcf7l1a^-/-^* embryos scored in each experiment. Eye formation was scored as rescued when pigmented retinal tissue was evident.

**Table S6 Size of the eye profile area is smaller in *mo^SPtcf7l2^* injected embryos at 30hpf.**

Volume in µm^3^ of the eye profile of 32hpf fixed embryos from wildtype embryos injected with *mo^C^* or *mo^SPtcf7l2^*. Avg, Average; SD, Standard Deviation.

**Table S7** Results from luciferase reporter assay experiments expressed in relative light units using *FLAG-Ax2* to induce Wnt activity. Avg, Average; SD, Standard Deviation; %, percentage relative to *FLAG-Ax2* condition.

**Table S8** Results from luciferase reporter assay experiments expressed in relative light units using *VP16-TCF7L2* to induce Wnt activity. Avg, Average; SD, Standard Deviation; %, percentage relative to *VP16-TCF7L2* condition.

**Table S9 Peptides recovered by mass spectrometry and their respective modifications.**

## Supplementary Materials and Methods

### Genotyping and adult tissue dissection

Genomic DNA was extracted from methanol or 4% paraformaldehyde fixed embryos by incubating in 25µl of KOH 1.25M, EDTA 10mM at 95°C for 30min, and then neutralised with 25µl of Tris-HCl 2M.

For tissue dissection, adult fish around a year old were culled by deep anesthetising in ice-cold 0.15% 2-phenotyethanol. Liver and pancreas tissue were isolated under a dissecting fluorescence microscope using *Tg*(*fabp10:dsRed*)*^gz4^* (Dong et al., 2007) and *Tg*(*XlEef1a1:GFP*)*^s854^* (Field et al., 2003) lines to distinguish liver and pancreas respectively. Other organs had clear anatomical boundaries and were easily dissected.

### Luciferase Reporter HEK cell transformation and assay conditions

HEK293 cells were grown routinely in DMEM + 10% foetal bovine serum and penicillin/streptomycin (all Gibco from Thermo-Fisher/Invitrogen). White-sided, clear-bottomed 96 well plates (Corning) were seeded at 8,000 cells/well in 100µl/well medium without antibiotics. All transfections were carried out with Transfectin (Bio-Rad) at a ratio of 3/1 (0.3 µl Transfectin/100ng DNA). Competition assays were carried out using VP16-TCF4 as the constitutive TCF (Ewan et al., 2010). The following plasmid amounts were used: VP16-TCF4: 43.5 ng/well; 25 ng/well reporter plasmids; 25 ng/well TCF isoform +pcDNA; 6.5 ng/well pcDNA-lacZ totalling 100 ng/well DNA. Transfectin was pipetted into 12.5 µl/well OptiMEM (Gibco/Thermo-Fisher) and the appropriate amount of DNA (above) into another 12.5 µl/well OptiMEM. The two were mixed and complexes were allowed to form over 20 min. The complexes were applied to the cells for 4 hrs after which the medium was removed and replaced with a fresh 100 µl/well medium. After 2 days incubation, the medium was removed the cells were lysed in 50 µl/well GLO Lysis buffer (Promega) over 20 min on slow shake. The lysate was then split into two 25 µl/well fractions. The luciferase assay was carried out with 25 µl/well Bright-GLO (Promega) and the lacZ transfection control assay with 25 µl/well Beta-GLO (Promega). All measurements were taken on a Fluostar plate reader (BMG) set to luminescence mode.

### HEK293 cell immunohistochemistry and co-immunoprecipitation protocol

For immuno-histochemistry, following cell fixation and permeabilisation, samples were washed with PBS, blocked in 5% albumin, and then incubated at 4°C overnight in mouse anti-MYC antibody high-affinity 9E10 (SCBT) diluted 1:1000 in 1% albumin in PBS. Cells were washed with PBS and incubated at room temperature for 1 h in Alexa Fluor 568 goat anti-mouse (Molecular Probes, Eugene, OR) diluted 1:1,000 in 1% albumin in PBS. Samples were washed with PBS, incubated in DAPI (1:50,000) for 5 min at room temperature, washed again, and mounted.

For Co-IP, following transformation, 350µl of Lysis Buffer 1 (below) was added and cells were resuspended off the plate by pipetting and lysed on ice for 1hr gently mixing every 10min. Samples were then centrifuged at 16.000g for 15min at 4°C, to enable removal of the supernatant and samples were then stored on ice. 70µl of Lysis Buffer 2 was added to the pellet which was then sonicated three times for 5 cycles at 40% output (Branson 450 sonicator). Tubes were centrifuged at 16.000g for 15min at 4°C and the supernatant was collected and added to the supernatant of the previous centrifugation step. Proteins were quantified by BCA method (Sigma, BCA1).

Lysis Buffer 1: NaCl 150mM, Tris-HCl 80mM pH7.2, NP-40 0.5%, Glycerol 20%. 10µl of complete protease Inhibitor (Sigma, P-8340), 10µl of phosphatase Inhibitor 1 (Sigma, P-2850) and 10µl of phosphatase Inhibitor 2 (Sigma, P-5726) were added before protein extraction for each 1ml of lysis buffer.

Lysis Buffer 2: NaCl 300mM, Tris-HCl 20mM pH7.2, SDS 0.01%, Triton 1%, EDTA 2mM, add 10µl of complete protease Inhibitor (Sigma, P-8340), 10µl of phosphatase Inhibitor 1 (Sigma, P-2850) and 10µl of phosphatase Inhibitor 2 (Sigma, P-5726) were added before protein extraction for each 1ml of lysis buffer.

For each Immunoprecipitation (IP) experiment condition 40µl of anti-myc bead slurry (Sigma, A7470) was added to 500µl of protein sample diluted to 1µg/µl in IPP buffer (Lysis Buffer 1 but with 5mM EDTA and no Glycerol), and incubated for 4hrs at 4°C in a small rotator mixer at 4RPM. Beads were spun for 30sec in a top bench centrifuge, the supernatant removed, and 500µl of IPP buffer added, and rinsed three times. Samples were centrifuged at 12000g for 1min at 4°C, the supernatant was removed and 40µl of 1.5x Laemmli Buffer was added. Proteins were eluted from the beads by incubating at 95°C for 5min.

### Yeast Transformation

Fresh Y2Gold yeast colony was used to inoculate 3ml of Yeast Peptone Dextrose Adenine (YPDA) liquid media for 8hrs and then 50µl of the culture was used to inoculate 50ml of YPDA and grow over night. The culture was grown until OD^600^ was 0.16 and then centrifuged at 700g for 5min at room temperature. The supernatant was discarded and the yeast resuspended in 100ml of fresh YPDA liquid media. The culture was grown for 3hrs or until OD^600^ reached 0.4-0.5, and centrifuged at 700g for 5min at room temperature, supernatant was discarded and the yeast pellet was resuspended in 60ml of water. Yeast were then centrifuged at 700g for 5min at room temperature, supernatant was discarded and yeast were resuspended in 3ml of 1.1X TE/LiAc (Tris-HCl 1mM, EDTA 1.1mM, Li Acetate 110mM, pH 7.5), centrifuged at 16000g for 15 sec, supernatant was discarded and yeast were resuspended in 1.2ml of 1.1 TE/LiAc. Competent yeast were kept at room temperature and used within less than an hour.

For yeast transformation, herring testis DNA (10µg/µl, Sigma, D-6898) was denatured at 100°C for 5min and then chilled on ice. Cells were transformed by mixing 50µg of herring testis DNA, 400ng of bait and prey DNA (80ng/µl), 50µl of competent yeast, 500µl of PEG/LiAc (40% Polyethilene glycol (Sigma, P-3640), Lithium Acetate 100mM), vortexing and incubating at 30°C for 30 min mixing every 10min. After this 20µl of DMSO were added, cells were mixed by inversion, and heat shocked at 42°C for 15min, mixing every 5min. Cells were then chilled on ice for 2min, centrifuged at 16000g for 10 sec, supernatant was discarded and yeast were resuspended in 200µl of sterile TE 1X. All the yeast were plated on specific dropout selective agar plates (Materials and Methods).

